# In the *Drosophila* germline H2Av and Arp6 suppress transposons by driving piRNA pathway expression

**DOI:** 10.64898/2026.05.19.726329

**Authors:** Norbert Andrási, Hannah M. Ryon, Yicheng Luo, Katalin Fejes Tóth

## Abstract

The spatiotemporal control of transcription and the maintenance of germline genome integrity depend on dynamic chromatin architecture. In *Drosophila*, the actin-related protein Arp6—a core subunit of the SWR1-like Domino chromatin remodeling complex—mediates the deposition of the histone variant H2Av. Previous studies have established H2Av as a key transcriptional regulator that modulates the +1 nucleosome barrier to promote RNA Polymerase II (Pol II) pause release and productive elongation. Conversely, H2Av is also integral to heterochromatin assembly and gene silencing. Here we demonstrate that Arp6 and H2Av are essential for female fertility and the global repression of transposable elements (TEs) in the *Drosophila* ovary. Rather than repressing TEs directly, we show that Arp6 and H2Av maintain genomic stability indirectly by driving the transcription of core PIWI-interacting RNA (piRNA) pathway genes. Depletion of either chromatin factor leads to a significant loss of piRNAs and reduced non-canonical transcription of dual-strand piRNA clusters. This defect stems from a failure to express the Rhino-Deadlock-Cutoff (RDC) complex, alongside the downregulation of multiple other piRNA biogenesis factors. Genomic profiling confirms that H2Av acts predominantly as an activating signal at host gene promoters. Upon H2Av or Arp6 depletion, genes that rely on H2Av for their expression exhibit a distinct upstream shift and more precise spatial localization of the Pol II peak at the TSS, indicating an impaired transition from transcription initiation into productive elongation. Together, our findings build upon the known transcriptional activation functions of the Arp6-H2Av axis, revealing that this established chromatin mechanism is critical for licensing piRNA-mediated genome defense and ensuring germline maintenance.

## Introduction

Eukaryotic gene expression is governed by the packaging of DNA into chromatin, which establishes a restrictive environment for DNA-templated processes (Struhl 1999, Weber et al. 2014). For many metazoan genes, the rate-limiting step in transcription is not the initial recruitment of Pol II, but rather the transition from a promoter-proximal paused state into productive elongation (Rougvie and Lis 1988, Rasmussen and Lis 1993, Adelman and Lis 2012, Weber et al. 2014). Following initiation, Pol II frequently halts 20 to 60 nucleotides downstream of the transcription start site (TSS), encountering the entry site of the first (+1) nucleosome (Kwak et al. 2013, Day et al. 2016). This +1 nucleosome represents a mechanical and energetic barrier to elongating polymerases (Dvir et al. 1996, Weber et al. 2014, Core and Adelman 2019). To overcome this physical obstacle and facilitate transcription, cells actively alter the biophysical properties of the +1 nucleosome by replacing canonical histones with specialized, non-canonical variants (Henikoff and Ahmad 2005, Saunders et al. 2006, Teves et al. 2014, Weber and Henikoff 2014, Li et al. 2023).

In *Drosophila melanogaster*, this critical function is fulfilled by the unique histone variant H2Av (Mavrich et al. 2008b, Weber et al. 2010, 2014). The targeted, replication-independent deposition of H2Av into chromatin is catalyzed by the SWR1-like DOMINO chromatin remodeling complex, DOM-B (Börner and Becker 2016, Scacchetti et al. 2020). Actin-related protein 6 (Arp6) is an essential core subunit of DOM-B required for the efficient exchange of canonical H2A for H2Av (Scacchetti et al. 2020). Consequently, Arp6 mutations closely mimic H2Av depletion phenotypes (Hsiao et al. 2023), making Arp6 a critical factor for investigating H2Av-dependent chromatin dynamics and transcriptional control. Once deposited, H2Av imparts a complex, context-dependent dual function on chromatin, participating in both the activation and repression of transcription (Swaminathan et al. 2005, Hanai et al. 2008, Kotova et al. 2011, Baldi and Becker 2013, Ibarra-Morales et al. 2021, Hsiao et al. 2023, Chen et al. 2025).

At actively transcribed genes, H2Av is heavily enriched at the +1 nucleosome, where structurally destabilizes the nucleosome and lower the energy barrier for Pol II, thereby facilitating the release of paused Pol II into productive elongation (Mavrich et al. 2008b, Weber et al. 2010, 2014). Loss of H2Av heightens this nucleosomal barrier, resulting in aberrant Pol II stalling and reduced transcriptional output (Weber et al. 2014). Paradoxically, beyond its role in transcriptional activation, *Drosophila* H2Av is also required for the establishment of constitutive heterochromatin and gene silencing (Swaminathan et al. 2005, Hanai et al. 2008, Baldi and Becker 2013, Giaimo et al. 2019). Its incorporation licenses the subsequent acetylation of H4K12, a prerequisite for H3K9 methylation and the recruitment of Heterochromatin Protein 1 (HP1) to repressive domains(Swaminathan et al. 2005).

How the Arp6-H2Av chromatin axis switches between facilitating active Pol II elongation and nucleating transcriptional silencing remains a central question in genome biology. This question is particularly paramount in the metazoan germline, where mobilization of transposable elements (TEs) is a source of continuous threat of genomic instability, necessitating their silencing (Bingham et al., 1982; Kidwell et al., 1977). To protect germline integrity, animals rely on an adaptable, small RNA-based immune system known as the PIWI-interacting RNA (piRNA) pathway, which utilizes small non-coding RNAs to guide Piwi-clade Argonaute proteins to silence TEs via post-transcriptional cleavage and co-transcriptional heterochromatin formation (Vagin et al. 2006, Brennecke et al. 2007, Gunawardane et al. 2007, Li et al. 2009, Malone et al. 2009, Sienski et al. 2012, Le Thomas et al. 2013, Rozhkov et al. 2013, Webster et al. 2015, Rogers et al. 2017, Ninova et al. 2020).. Large-scale, genome-wide RNAi screens previously identified both H2Av and Arp6 as factors required for TE silencing in the *Drosophila* ovary (Czech et al. 2013, Handler et al. 2013). However, given the dual nature of H2Av, it has remained unclear whether the Arp6-H2Av axis regulates TEs directly—by facilitating heterochromatin assembly at individual TE loci—or indirectly, possibly by modulating the piRNA pathway by a yet-to-be-identified mechanism.

In this study, we dissect the molecular role of Arp6 and H2Av in TE silencing and transcriptional control within the *Drosophila* female germline. We demonstrate that Arp6 and H2Av are indispensable for female fertility and the global repression of transposons. Rather than acting directly at TE loci, our genomic and transcriptomic analyses reveal that the Arp6-H2Av axis serves primarily as an activating signal at host gene promoters to drive the transcription of essential piRNA pathway components. Depletion of Arp6 or H2Av precipitates a severe collapse of piRNA cluster transcription and subsequent piRNA loss. Furthermore, precise mapping of engaged Pol II demonstrates that H2Av and Arp6 depletion alters promoter-proximal pausing at target genes. Downregulated genes exhibit an upstream shift and narrowing of the Pol II peak, signifying an impaired ability to traverse the +1-nucleosome barrier and transition into productive elongation. Together, our results uncover an essential, indirect mechanism of TE control whereby the H2Av and the Dom-B complex ensure genome integrity by licensing the transcription of the piRNA pathway itself.

## Results

### H2Av and Arp6 are required for female germline development and global transposon repression in the fly ovary

Genome-wide studies have identified multiple factors critical for transposon repression (Czech et al. 2013, Handler et al. 2013, Muerdter et al. 2013). While the mechanisms of action for many piRNA pathway components have been well established (Iwasaki et al. 2015, Huang et al. 2017, Ozata et al. 2019), the molecular functions of other identified factors remain unresolved. Here, we focus on two such factors: the histone variant H2Av and the Actin-Related Protein 6 (Arp6), a core component of the SWR1 complex responsible for swapping the canonical H2A histone with H2Av. To investigate their function, we depleted these genes specifically in the ovarian germline by crossing flies expressing the *nanos-GAL4-VP16* driver to those carrying UASp-driven short hairpin RNAs (shRNAs). Because publicly available *Arp6* hairpins yielded insufficient depletion, we generated new transgenic hairpin lines. RT-qPCR validation confirmed robust knockdown (KD) efficiencies, showing an 85% reduction in *arp6* and a 90% reduction in *H2Av* transcript levels (normalized to the geometric mean of two reference genes, *rp49* and *gapdh1*), compared to control flies expressing an shRNA against the *white* gene (shW) (Fig. S1A).

To confirm previous observations from genome-wide screens indicating that H2Av and Arp6 impact female fertility (Czech et al. 2013, Handler et al. 2013), we crossed KD females to wild-type males and quantified eclosing progeny. Germline knockdown (GLKD) of either *Arp6* or *H2Av* resulted in complete female sterility, yielding zero eclosed flies compared to an average of 107 (SD = 4.8) in the shW control (Fig. 1A). This requirement appears sex-specific; when we depleted *arp6* or *H2Av* in the male germline using the same *nanos-GAL4-VP16* driver, we observed only a slight decrease in hatching rates (Fig. S1B). *H2Av* KD ovaries were noticeably diminished in size. *Arp6* KD also led to reduced ovarian size, though with variable penetrance, which likely reflects the slightly weaker knockdown efficiency achieved for this gene (Fig. 1B).

**Figure 1.**
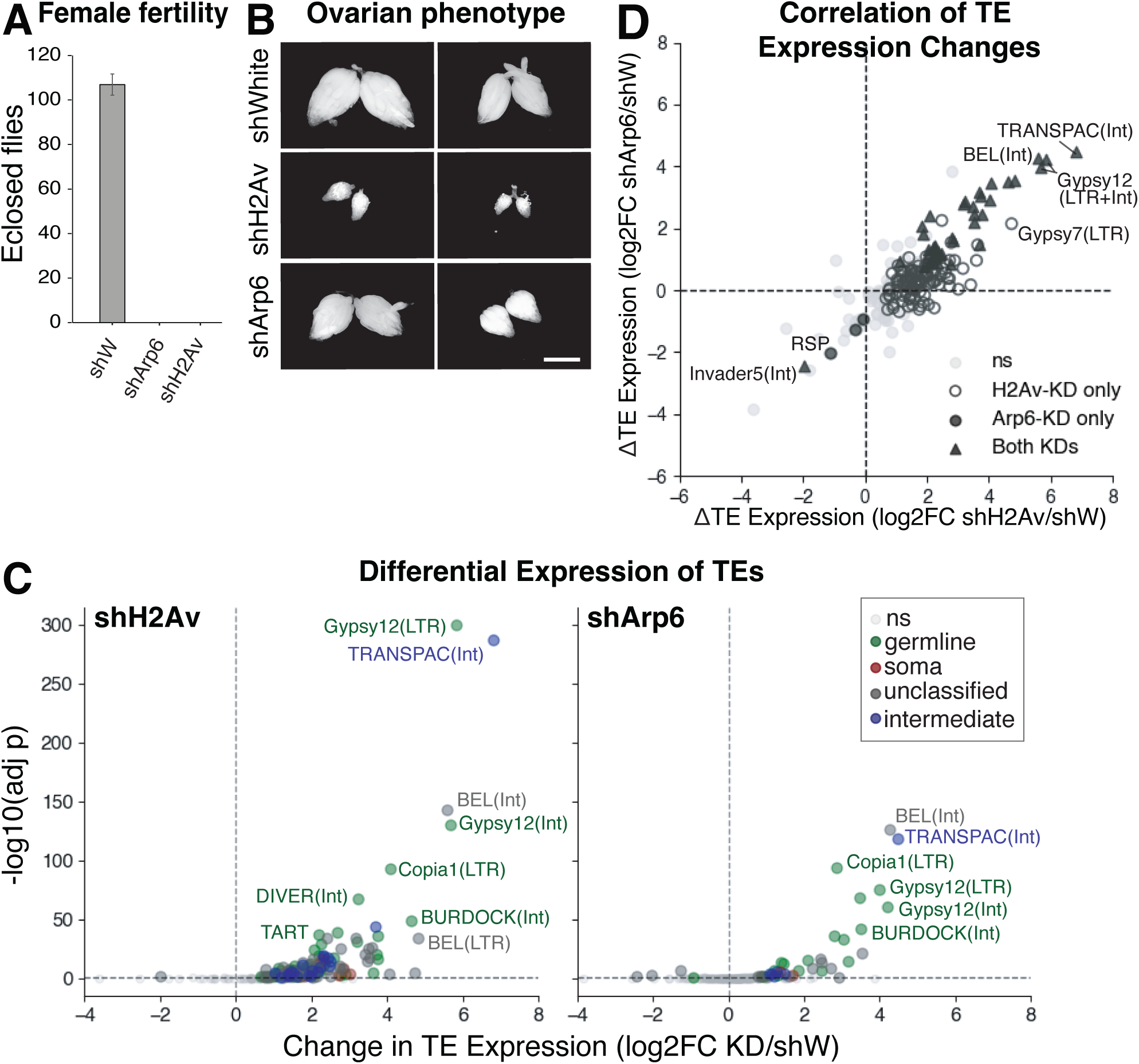
His2Av and Arp6 are required for normal germline development and transposon silencing. (A) Loss of H2Av and Arp6 leads to female infertility. Compromised fertility of H2Av and Arp6 GLKD compared to white GLKD females (nos-Gal4-VP16 driver was used, error bars represent SD). (B) Loss of H2Av and Arp6 impairs oogenesis and leads to smaller ovaries. Ovary phenotype of white, H2Av and Arp6 GLKD flies (Scale bar: 0.5 mm). (C) H2Av (left) and Arp6 (right) are required for global TE repression in the female germline. TE families are colored based on their somatic, germline, intermediate, or unclassified tissue-specific expression. Ns: non-significant change (D) H2Av and Arp6 regulate the same TEs. Differential expression of all TEs (TEtranscript with DESeq2-based statistical analysis) with significant points for each condition, or shared in both conditions, highlighted using different markers.

Because atrophied ovaries and sterility are hallmark phenotypes of TE dysregulation (Schüpbach and Wieschaus 1991, Cox et al. 1998, Klattenhoff et al. 2007, Li et al. 2009, Siomi et al. 2011, Sienski et al. 2012), we next performed global RNA-seq on ovaries from control and GLKD flies to assess transposon expression. Differential expression analysis revealed a global upregulation of TEs upon loss of either gene. In alignment with our germline-specific driver, this predominantly affected germline-specific TEs. Of the 217 annotated TE families, 142 were significantly upregulated in the *H2Av* GLKD, and 37 of these were concurrently upregulated upon *Arp6* depletion (Fig. 1C). The magnitude and significance of TE derepression were notably greater in the *H2Av* KD, mirroring its more severe morphological phenotype. We further validated the upregulation of select TE families via RT-qPCR (Fig. S1C).

Given that Arp6 is essential for the SWR1-mediated deposition of H2Av (Scacchetti et al. 2020), we hypothesized that disrupting either factor would yield overlapping transcriptional defects. Indeed, we observed a highly significant correlation (Spearman’s ρ = 0.705, p = 2.9 × 10⁻²³) in the expression changes of affected TEs between the two KDs (Fig. 1D). Taken together, these data demonstrate that H2Av and the SWR1 complex act within the same functional pathway to silence TEs, maintain normal ovarian morphology, and ensure female fertility.

### TEs are repressed by H2Av mostly by a piRNA-mediated mechanism

Given H2Av’s known nuclear transcriptional functions (Swaminathan et al. 2005, Kotova et al. 2011, Baldi and Becker 2013, Ibarra-Morales et al. 2021, Chen et al. 2025), we first investigated whether it represses TEs through direct transcriptional regulation. Using ChIP-seq with a validated pan-H2Av antibody, we quantified H2Av enrichment over TE promoters (TSS +/- 1kb). If H2Av directly represses these loci, it should be enriched at TEs—particularly those upregulated upon its depletion. However, for most TE families, we observed neither significant H2Av enrichment nor a correlation between wild-type H2Av signal intensity and the degree of derepression upon H2Av KD (Fig. 2A, Fig. S2A). Notably, among the few derepressed TEs that did display H2Av enrichment in the shW control, the highest signals were observed at the telomeric TEs: *HetA*, *TART*, and *TAHRE*.

**Figure 2.**
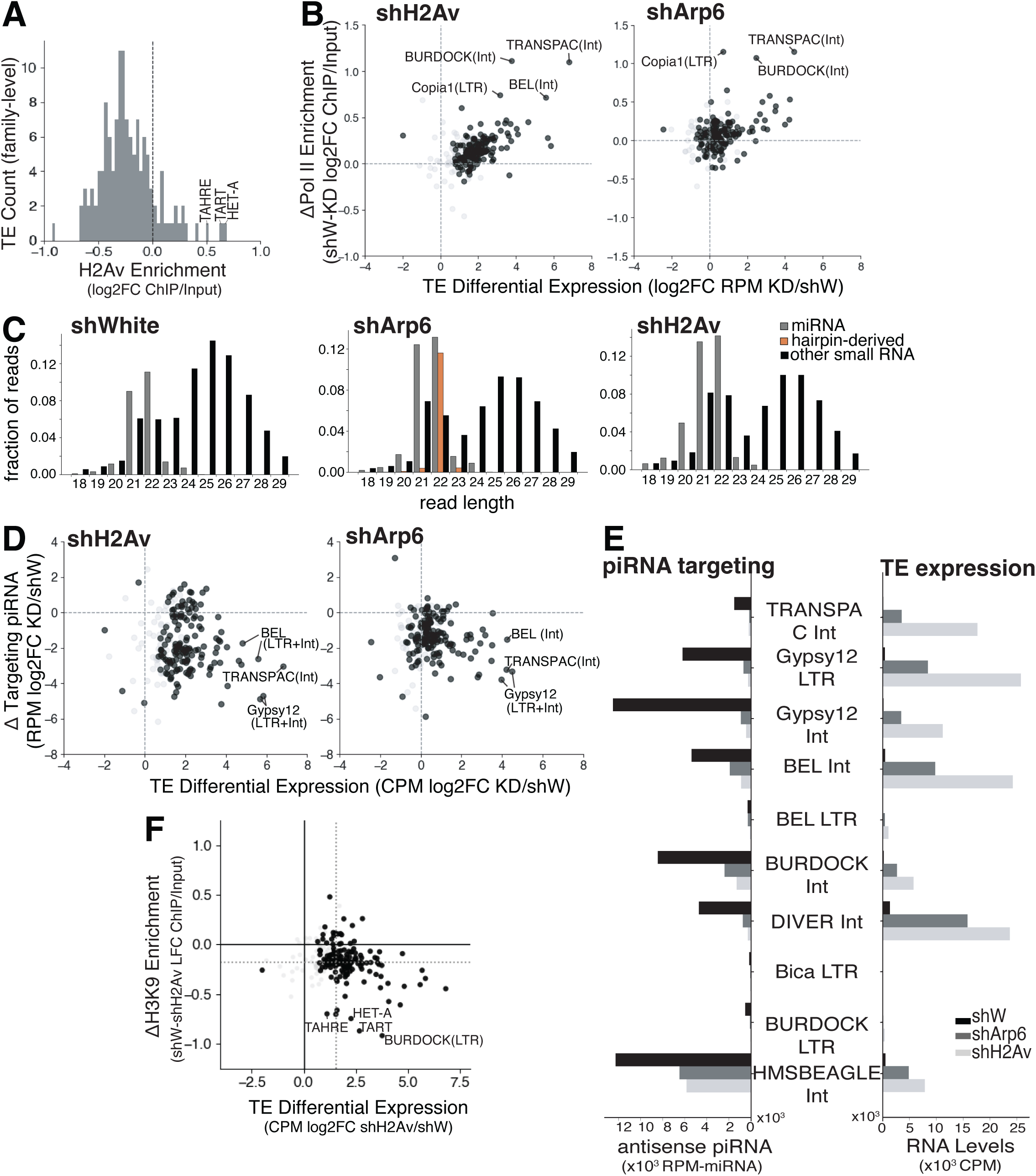
TE derepression in the absence of H2Av and Arp6 is primarily due to the loss of silencing piRNAs. (A) H2Av is only enriched at telomeric TEs. H2Av signal distribution at the promoter regions (+/-1kb TSS) of TEs (aggregated by family and averaged for 2 reps). (B) TE transcript expression changes weakly correlate with Pol II accumulation changes upon H2Av/Arp6 GLKD. Change in Pol II ChIP signal enrichment vs change in TE expression upon H2Av GLKD (left) and Arp6 GLKD (right) relative to shW control analyzed at the family level. (C) Arp6 and H2Av GLKD leads to loss of piRNAs. Small RNAs were separated into miRNAs and other small RNAs, based on miRbase sequence alignment, and the average length distribution plotted (2 replicates). shRNA-derived sequences were plotted separately. (D) TE upregulation correlates with loss of antisense “targeting” piRNA upon H2Av or Arp6 GLKD. Scatter plot showing the relationship between changes in piRNA targeting and TE expression upon His2Av (left) and Arp6 (right) knockdowns; (E) TE expression (right) and corresponding piRNA targeting (left) in shW (black), shArp6 (dark grey), shH2Av (light grey) for the top 10 most derepressed TEs in H2Av GLKD. (F) TE derepression is not due to H3K9me3 loss on most TEs. Change in H3K9me3 levels across the body of all TEs vs change in TE expression in the H2Av GLKD compared to White GLKD, with solid lines showing 0, and dashed lines show the population median for either axis. In all scatterplots (B, D, and F), significantly derepressed TE families (padj<0.05) are shown in black (nonsignificant in grey) with significance defined based on H2Av GLKD for both conditions.

Despite the general lack of H2Av accumulation at TEs, we next asked whether TE derepression occurs at the transcriptional level. ChIP-seq for initiating RNA Polymerase II (Pol II pSer5) revealed that, although baseline Pol II signal is below input for most TEs in the control, there is a modest but significant global increase in Pol II enrichment at the promoters of derepressed TEs upon H2Av depletion (median Δ = 0.15, Wilcoxon p = 7.3 × 10⁻²¹), and to a lesser extent upon Arp6 depletion (median Δ = 0.08, Wilcoxon p = 4.7 × 10⁻²¹) (Fig. 2B, S2B). Together, the lack of H2Av occupancy combined with increased Pol II recruitment indicates that TE activation upon the loss of H2Av is primarily an indirect consequence of lost transcriptional repression, with telomeric TEs serving as a notable exception of potential direct regulation.

Since the piRNA pathway is the primary mode of TE repression in the *Drosophila* ovary (Vagin et al. 2006, Brennecke et al. 2007, Malone et al. 2009), we explored whether this global TE upregulation resulted from deficient piRNA silencing. Small RNA-seq of GLKD and control ovaries revealed a sharp drop in the piRNA-to-miRNA ratio—falling from 2.74 in the shW control to 1.24 and 1.47 in shH2Av and shArp6 samples, respectively (Fig. 2C), indicating a global reduction of the piRNA pool. To determine if this reduction drove TE derepression, we correlated changes in targeting piRNAs with changes in TE expression. We observed a marked global shift: the increased expression of most TE families was tightly coupled with the loss of their corresponding antisense piRNAs (Arp6 KD: median projection = 1.19, p = 2.2 × 10⁻³⁰; H2Av KD: 2.43, p = 1.3 × 10⁻³⁴) (Fig. 2D). This was especially pronounced for the 10 most upregulated TEs (Fig. 2E); for instance, the *gypsy12* retroelement increased 51.2- and 15.6-fold in H2Av and Arp6 GLKDs, respectively, while its targeting piRNAs plummeted 29.5- and 13.8-fold. Overall, 72% of upregulated TEs in shArp6 and 66% in shH2Av can be directly attributed to the loss of piRNAs (LFC < -1), identifying deficient piRNA-mediated silencing as the primary driver of TE activation.

To detect evidence of impaired piRNA-mediated transcriptional silencing, we assessed H3K9me3, a repressive histone modification guided to TEs by the piRNA pathway (Sienski et al. 2012, Le Thomas et al. 2013, Rozhkov et al. 2013). Family-level quantification of H3K9me3 ChIP signal confirmed there is a modest, but significant, global decrease in H3K9me3 occupancy at TEs, and this loss correlates with increased TE expression in the H2Av KD (p = 1.5 × 10⁻²⁷) (Fig. 2F). While only a small subset of TEs underwent substantial H3K9me3 occupancy changes in the H2Av knockdown, the three telomeric TEs (*HetA*, *TART*, and *TAHRE*) were notably among them (Fig. 2F). These results suggest that piRNA-mediated H3K9me3 deposition is impaired upon H2Av loss.

While the majority of TEs are derepressed due to piRNA loss, approximately 30% of activated TEs (50 families) could not be confidently attributed to this mechanism (targeting piRNA LFC ≥ -1). This prompted us to revisit the possibility of direct H2Av regulation as an alternate, albeit rare, silencing mechanism. Across the genome, our ChIP-seq analysis identified only 21 TE families with H2Av enrichment (ChIP signal > input) in wild-type ovaries. Because 15 of these 21 families also experienced a substantial loss of targeting piRNAs upon H2Av KD (LFC < -1), it is difficult to uncouple direct H2Av regulation from indirect piRNA-mediated effects at those loci. However, the remaining 6 H2Av-enriched TEs were significantly upregulated despite little-to-no loss of their targeting piRNAs (Fig. S2C). Notably, three of these six were the telomeric transposons (*HetA*, *TART*, and *TAHRE*), which displayed the strongest H2Av enrichment (Fig. 2A, S2A) and some of the most drastically decreased H3K9me3 levels in shH2Av (Fig. 2F, S2C). These observations support a two-pronged TE-silencing function for H2Av: the vast majority of TEs are regulated indirectly via maintenance of the piRNA pathway, while a small subset of TEs—most prominently at the telomeres—are regulated through a separate, likely direct and chromatin-based, mechanism.

### H2Av is required for piRNA production and cluster expression

H2Av and Arp6 could regulate piRNA biogenesis through distinct mechanisms. They might facilitate the establishment of a cluster-specific chromatin state essential for piRNA precursor transcription (e.g., at dual-strand clusters like *42AB*). Alternatively, they could act independently of cluster transcription, regulating downstream precursor processing via the ping-pong amplification loop (Brennecke et al. 2007, Gunawardane et al. 2007, Czech and Hannon 2016) or the phased processing pathway (Han et al. 2015, Mohn et al. 2015).

To determine whether H2Av or Arp6 depletion impacts ping-pong amplification, we analyzed the characteristic 10-nt overlap between the 5’ ends of complementary piRNAs mapping to TE sequences (Brennecke et al. 2007, Gunawardane et al. 2007). This ping-pong signature remained unaltered upon Arp6 GLKD and decreased only moderately upon H2Av GLKD, with the average Z_10_-score dropping from 10.9 in the shW control to 6.8 in shH2Av ovaries (Fig. 3A). We next examined the phased piRNA biogenesis pathway, wherein Piwi binds precursor transcripts at 5’-cleavage sites (generated by ping-pong processing) followed by phased transcript cleavage by the endonuclease Zucchini (Han et al. 2015, Mohn et al. 2015). Quantification of the phasing signature revealed the characteristic 1-nt bias across all three samples (Z1-score shW: 7.79, shArp6: 7.48, shH2Av: 7.92), showing no evidence of disrupted phasing in the knockdowns (Fig. 3B). Thus, while H2Av moderately influences the ping-pong cycle, processing defects alone cannot explain the profound global reduction in mature piRNAs.

**Figure 3.**
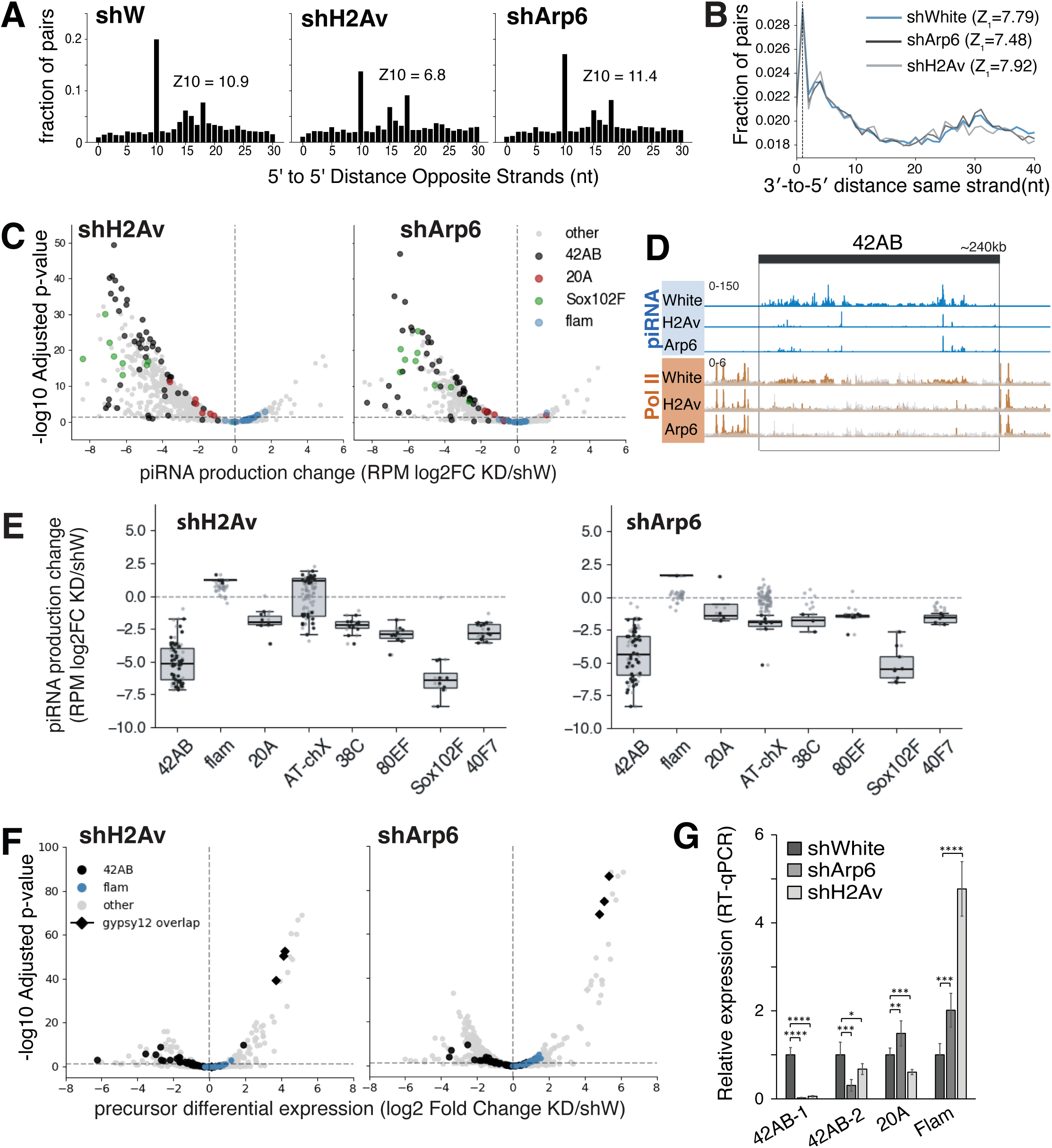
Loss of Arp6 and H2Av result in decreased cluster transcription without major disruption to ping-pong and phased piRNA processing. (A) Ping-pong amplification is only mildly reduced by germline KD of H2Av and unaffected by loss of Arp6. (B) Phasing signature is unchanged upon GLKD of H2Av and Arp6. Shown is the distribution of distances between the 5’ end of one piRNA and the 3‘end of the nearest upstream piRNA with characteristic 1nt enrichment. (C) piRNAs are lost from several germline clusters. 2 replicate average fold change in piRNA coverage over 5kb windows (unique mappers only) normalized to miRNA counts was plotted. (D) IGV browser view of piRNA uniquely mapping at 42AB and flamenco showing library-size normalized bigwig tracks for a single representative replicate. (E) Strip plots of differential piRNA production upon H2Av (left) and Arp6 (right) GLKD over 5kb windows in the main piRNA-producing clusters compared to shW and normalized by miRNA. Significant points shown in black, nonsignificant in grey. (F) piRNA cluster precursor expression (mRNA levels) shows a more complex response to knockdowns with both increases and decreases observed amongst the same 5kb windows. Windows overlapping gypsy12 are shown with a diamond marker. (G) His2Av and Arp6 GLKD leads to loss of germline cluster precursors. RT-qPCR showing changes in piRNA precursor expression (normalized to the geometric mean expression of *RP49* and *GAPDH1*) at selected piRNA clusters (Student’s t-test, *p<0.05, **p<0.01, ***p<0.001, ****p<0.0001, error bars represent SD).

Because small RNA processing remains largely intact, we investigated whether piRNA production is compromised at the level of cluster transcription. Mature piRNAs are generated from long, single-stranded precursors originating from specialized genomic loci called piRNA clusters (Brennecke et al. 2007). In the *Drosophila* ovary, most germline piRNA clusters are transcribed bidirectionally by a non-canonical transcription mechanism (Li et al. 2009, Malone et al. 2009, Rangan et al. 2011, Chen et al. 2016, Hur et al. 2016). This process relies on the presence of the repressive H3K9me3 mark and the unique, cluster-specific Rhino-Deadlock-Cutoff (RDC) complex (Klattenhoff et al. 2009, Pane et al. 2011, Mohn et al. 2014, Zhang et al. 2014, Akkouche et al. 2017, Andersen et al. 2017, Chen et al. 2021). In contrast, piRNAs produced in the surrounding somatic follicular cells predominantly derive from the *Flamenco* cluster, a uni-stranded transcript generated by the canonical Pol II machinery (Prud’homme et al. 1995, Sarot et al. 2004, Brennecke et al. 2007, Mével-Ninio et al. 2007, Li et al. 2009, Malone et al. 2009, Goriaux et al. 2014). To understand how germline depletion of H2Av and Arp6 impacts these distinct loci, we mapped reads uniquely to each strand and quantified total piRNA abundance per cluster. To capture local intra-cluster variation, we also performed a windowed analysis using 5-kb non-overlapping bins.

We observed a significant decrease in mature piRNA production at most characterized germline piRNA clusters in the absence of H2Av and Arp6, with the most dramatic changes occurring at *42AB* and *Sox102* (Fig. 3C-E, S3A). The only exception was the *AT-chX* germline cluster, which showed little-to-no change. Conversely, piRNAs derived from the somatic uni-strand cluster, *Flamenco*, exhibited a significant increase in both total net production and window-based signal (Fig. 3C, E, S3A). This increase, particularly apparent upon H2Av depletion, is likely an artifact of the relative loss of germline tissue and the concomitant overrepresentation of follicular cells in the highly atrophied KD ovaries (Fig. 1B). Overall, mature piRNA production from major germline clusters is severely impacted by the depletion of H2Av and Arp6, corroborating the observed TE derepression.

To definitively evaluate whether these changes in mature piRNAs stem from altered precursor transcript levels, we performed RNA-seq on rRNA-depleted ovarian RNA. We mapped unique reads to the previously described 5-kb windows and aggregated the counts per cluster. Consistent with the loss of mature piRNAs, we detected a stark loss of long precursor transcripts across several dual-strand germline clusters. Exceptions included *38C* and *AT-chX*—which showed slightly increased median precursor expression per 5-kb window in both conditions—and *80EF*, which exhibited increased expression only in the absence of H2Av. These increases reflect activation of internal TE promoters (*copia, baggins1* and *gypsy 8* fragments, respectively), with read counts over all other windows within the clusters remaining very low (Fig. 3F, S3B). Similarly, the elevated mature piRNA signal at *Flamenco* was accompanied by an increase in precursor levels, further supporting the germline-to-soma ratio shift. These changes in precursor transcript levels demonstrated consistent correlation between the H2Av and Arp6 GLKDs (Fig. S3C) and were further validated via RT-qPCR of select clusters (Fig. 3G) and demonstrated consistent correlation between the H2Av and Arp6 GLKDs (Fig. S3C). Notably, a large RNA-seq peak appeared within *42AB* in the KD samples. This peak corresponds to the activation of an autonomously expressed *gypsy12* transposon integrated within the *42AB* locus, consistent with the loss of targeting piRNAs and the aberrant initiation of canonical Pol II transcription from within the cluster itself (Fig. 3F; Fig. S3B & D).

To confirm whether the loss of precursor transcripts is rooted in a failure of transcription initiation, we mapped Pol II pSer5 ChIP-seq data to cluster loci (allowing up to 50 multi-mappers, fractionally assigned to 5 kb tiles) (Baumgartner et al. 2022). Because Pol II pSer5 signal is intrinsically low across most wild-type piRNA clusters, quantitative tile-based analysis was largely uninformative and frequently confounded by sharp, but still relatively low, increases in Pol II signal at internal, derepressed TEs in the knockdowns (Fig. S3D). For example, cluster *80EF* showed a significant total increase in Pol II, but this localized entirely to the TSS of an internal *Gypsy8* element. Similarly, while the overall Pol II signal at *42AB* appeared weak or near zero in the tile-based scatterplots due to signal averaging over diffuse, noisy regions, locus-specific inspection revealed a different dynamic. In regions where piRNA mapping density was high in the shW control, we observed a broad, low-level Pol II occupancy (Fig. 3D). This broad Pol II signal was visibly reduced in both Arp6 and H2Av knockdowns, consistent with the loss of piRNA expression from these regions.

Finally, to rule out the possibility that H2Av directly regulates the transcription of these piRNA clusters, we mapped our H2Av ChIP-seq data to the cluster loci. We found no evidence of H2Av occupancy, with the sole exception of *40F7*, which exhibited strong H2Av enrichment exclusively at tiles overlapping an annotated gene promoter at one boundary of the cluster (Fig. S3E). The general lack of physical H2Av enrichment at piRNA clusters demonstrates that the transcription of piRNA precursors relies on Arp6 and H2Av indirectly. Overall, our findings reveal that H2Av and Arp6 maintain piRNA populations predominantly by licensing piRNA precursor transcription, functioning through an essential, indirect regulatory mechanism rather than direct chromatin binding at the clusters themselves.

### H2Av and Arp6 are critical for Rhino-Deadlock-Cutoff silencing complex activity

The non-canonical transcription of dual-strand germline piRNA clusters relies on the Rhino-Deadlock-Cutoff (RDC) complex, which recognizes repressive H3K9me3 marks and licenses long read-through transcription (Klattenhoff et al. 2009, Pane et al. 2011, Mohn et al. 2014, Zhang et al. 2014, Akkouche et al. 2017, Andersen et al. 2017, Chen et al. 2021). To determine if the loss of piRNA precursors in our knockdowns stems from disrupted cluster chromatin, we performed ChIP-seq for H3K9me3 and Rhino. Using the same 5 kb genomic tiling approach described for Pol II, we quantified levels of H3K9me3 and Rhino enrichment and observed a severe global depletion of Rhino in both knockdown conditions (Fig. S4A-B). This naturally results in the loss of Rhino occupancy at the clusters, with a particularly stark loss upon H2Av GLKD (Fig. 4A-B; Fig. S4A-C), and a strong correlation between the two KDs (Fig. S4D). Importantly, H3K9me3 levels at these loci remained largely unchanged or were only moderately reduced (Fig. 4A-B, Fig. S4B-C). Thus, the failure of Rhino to accumulate at clusters cannot be explained by the loss of its H3K9me3 binding sites.

**Figure 4.**
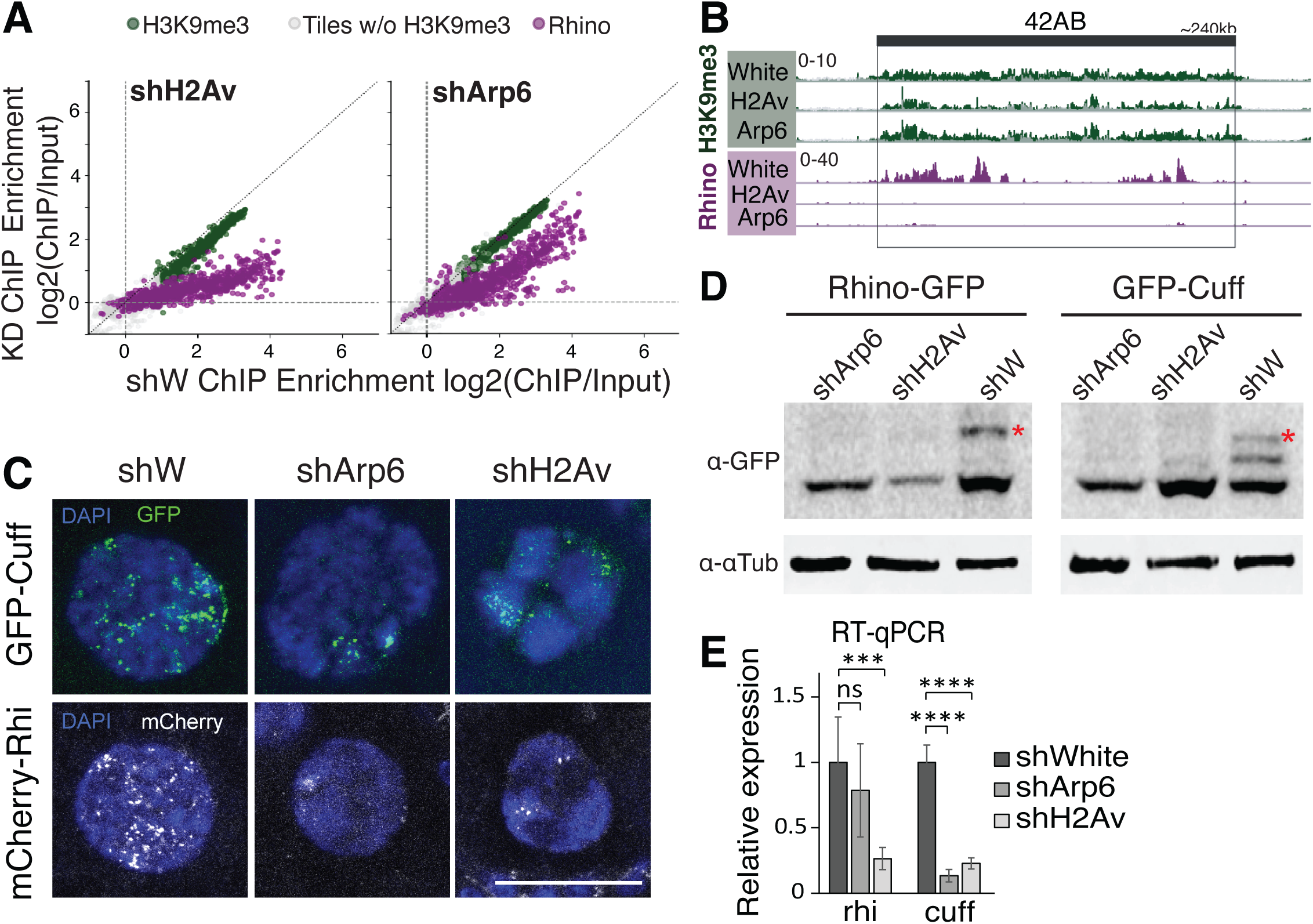
piRNA cluster transcriptional decrease occurs due to loss of RDC activity. (A) Rhi, but not H3K9me3 is lost over clusters upon H2Av or Arp6 GLKD. H3K9me3 and Rhino ChIP signal enrichment in control (shW) vs shH2Av (left) and shArp6 (right) over high mappability 5kb genomic tiles overlapping cluster annotations. (B) Representative single-replicate browser track viewed in IGV, showing 42AB with piRNA reads shown in blue, Pol II-Ser5 in orange, H3K9me3 IP in green, and Rhino IP in purple. ChIP input is shown in grey at same scale overlayed over the IP signal. Signal in input overlayed in gray. (C) Cuff and Rhi germline nuclear foci are lost upon H2Av and Arp6 GLKD. Confocal images of nurse cells nuclei showing GFP-Cuff and mCherry-Rhi localization (Scale bar: 20 µm). (D) Cuff and Rhi proteins are depleted from ovaries upon H2Av and Arp6 GLKD. Western blot analysis of Rhino-GFP and GFP-Cutoff protein from dissected ovaries. Tagged proteins were expressed under their respective endogenous promoter. Red asterix indicates the target protein. The lower band in Cuff-GFP might be degradation fragment of Cuff-GFP. (E) RT-qPCR showing normalized *rhi* and *cuff* mRNA levels normalized to the geometric mean expression of *RP49* and *GAPDH1* (Student’s t-test, ***p<0.001, ****p<0.0001, ns – not significant, error bars represent SD).

To further validate this loss of Rhino and test whether the entire RDC complex is destabilized, we visualized protein localization in nurse cell nuclei using transgenic flies expressing fluorescently tagged Rhi (mCherry-Rhi) or Cutoff (GFP-Cuff) under their native promoters. In both Arp6 and H2Av GLKDs, we observed a severe loss of Rhi and Cuff from nuclear foci (Fig. 4C). To distinguish between a failure of chromatin recruitment and an overall depletion of the proteins themselves, we performed Western blot analysis on ovarian lysates. Upon KD of either factor, both Cuff and Rhi proteins became completely undetectable (Fig. 4D), confirming a global loss of RDC proteins from germ cells.

Finally, we investigated whether this drastic protein loss originated at the transcriptional level. RT-qPCR analysis revealed a severe 7- and 4-fold reduction in *cutoff* (*cuff*) mRNA in the H2Av and Arp6 KDs, respectively (Fig. 4E). In contrast, *rhino* (*rhi*) mRNA levels were only marginally reduced in shArp6 and decreased ∼3.5-fold in shH2Av. To ensure these expression changes were not merely a consequence of germ cell loss in the atrophied KD ovaries, we performed RNA *in situ* hybridization. Consistent with the qPCR data, *rhi* transcripts remained visibly detectable, whereas *cuff* mRNA was essentially abolished upon depletion of either H2Av or Arp6 (Fig. S4E). Taken together, these findings demonstrate that Arp6 and H2Av are required to maintain *cuff* transcript levels. The collapse of *cuff* expression subsequently destabilizes the RDC complex (Mohn et al. 2014, Chen et al. 2021), driving the secondary degradation of Rhino protein, the failure of cluster transcription, and the ultimate derepression of TEs.

### H2Av and Arp6 depletion leads to global changes in ovarian gene expression, with enrichment of piRNA genes among those downregulated

To determine if the loss of *cutoff* reflects a broader collapse of the piRNA pathway and to evaluate the genome-wide impact of H2Av and Arp6 GLKD, we performed differential expression analysis of host genes using our ovarian RNA-seq datasets. Of the ∼14,000 annotated *Drosophila* genes, 1,993 (14.2%) were significantly upregulated and 1,700 (12.2%) were downregulated upon H2Av GLKD (|LFC| > 0.58, p < 0.05). Similarly, in the Arp6 GLKD, 546 (3.9%) and 429 (3.1%) genes were up-and downregulated, respectively. Comparing the two conditions revealed 479 upregulated and 321 downregulated genes in common (Fig. 5A). Consistent with a coordinated genome-wide function of these factors (Scacchetti et al. 2020, Ibarra-Morales et al. 2021, Hsiao et al. 2023, Chen et al. 2025), we observed a strong correlation (Spearman ρ = 0.73, p < 2.2 × 10⁻¹⁶) in the expression changes of host genes affected by the depletion of both proteins (Fig. S5A).

**Figure 5.**
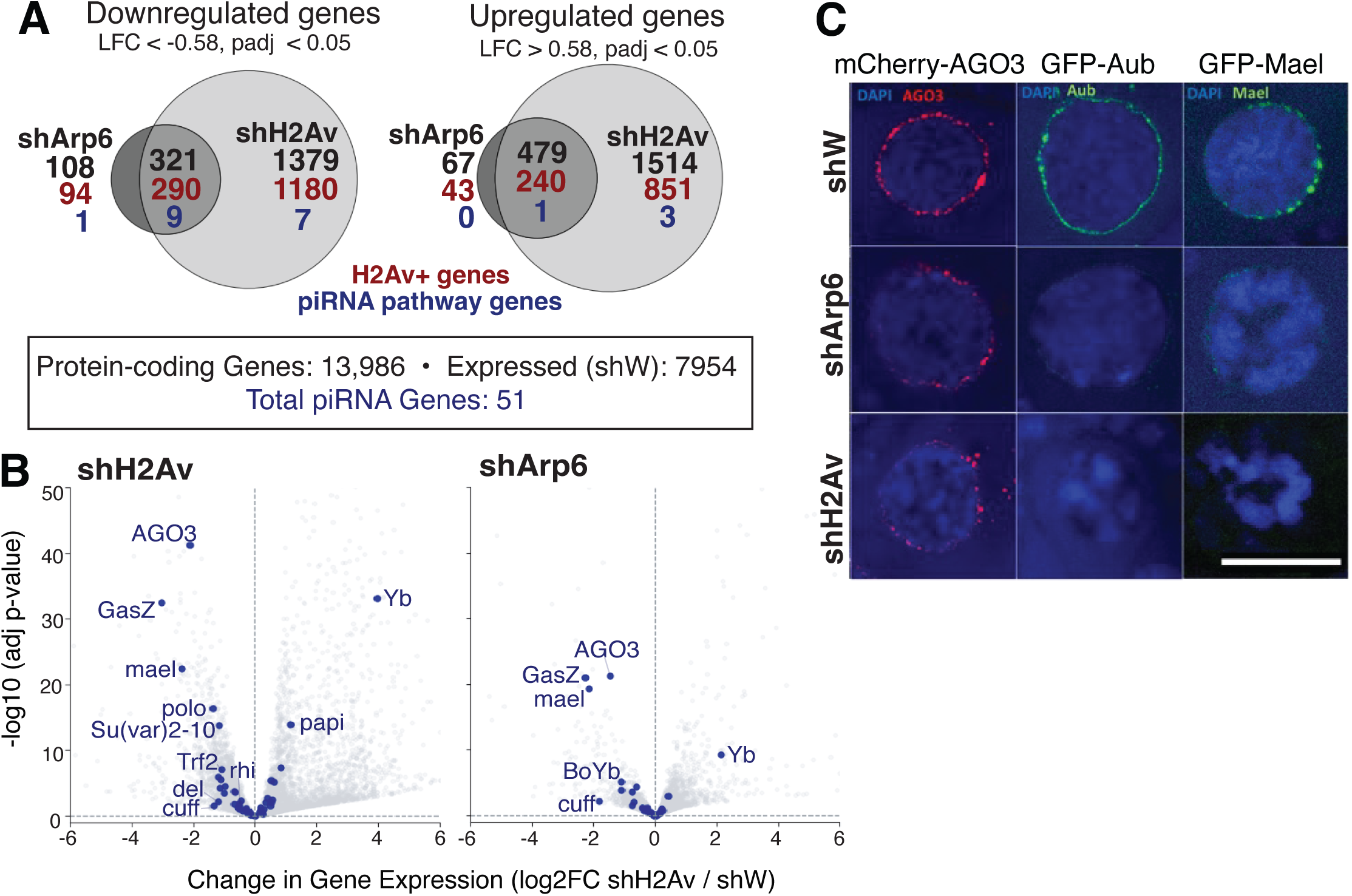
H2Av and Arp6 depletion leads to global changes in ovarian gene expression, with enrichment for piRNA genes among those downregulated. (A) Venn diagram of differentially regulated genes (padj <0.05, |LFC| > 0.58). Gene counts are shown in black for differential genes in shH2Av, shArp6, and both. Count of differential genes with H2Av signal (>0.4 LFC IP/Chip) is shown in red, and the count of differential genes involved in the piRNA pathway is shown in blue. (B) Most piRNA pathway genes depend on H2Av and Arp6 for their expression. RNA-Seq LFC and adjusted p-value based on DESeq2 differential analysis comparing the shH2Av (left) or shArp6 (right) to shW. piRNA-related genes are shown in blue. (C) AGO3, Aub and Mael expression is lost or greatly reduced in His2Av and Arp6 GLKD germline cells. Confocal images of nurse cells nuclei from flies expressing the respective tagged transgenic proteins driven by their endogenous promoter. (Scale bar: 20 µm).

Interestingly, while the overall number of up- and downregulated genes was relatively balanced in both knockdowns, focusing specifically on known piRNA pathway genes revealed a striking directional bias. Of the 51 established piRNA genes, 16 (31.4%) were downregulated in the H2Av GLKD and 10 (19.6%) in the Arp6 GLKD, with 9 of these core factors downregulated in both conditions (Fig. 5A-B). To test the significance of this overlap, we compared our data against a null distribution generated from 10,000 random gene sets of equal size. This revealed that piRNA genes are significantly enriched among downregulated targets in shArp6 (4-fold over expected, p = 0.0003). A weaker but still significant enrichment was observed in shH2Av (1.7-fold, p = 0.017), likely diluted mathematically by the much larger total number of downregulated genes in that condition. In contrast, only four piRNA genes (<7.8%) were upregulated upon H2Av depletion, and only one (*Yb*) upon Arp6 depletion. The upregulation of *Yb* in both KDs likely reflects an overrepresentation of somatic cells in the atrophied ovaries rather than a direct regulatory effect.

Among the most significantly downregulated genes in both KDs were central piRNA biogenesis and effector proteins, including *argonaute3*, *gasz*, and *maelstrom* (Fig. 5B; Fig. S5A). Consistent with our previous IF and qPCR results (Fig. S5B) and the loss of RDC, we also observed a statistically significant loss of *cutoff* and *deadlock* transcripts in both KDs, whereas *rhino* loss was significant only upon H2Av depletion (Fig. 5B). Additional factors such as *daed*, *BoYb*, and *trf2* were significantly downregulated in both KDs, while *su(var)2-10*, *egg*, *krimp*, *vret*, and *squ* showed significant loss exclusively upon H2Av depletion. Notably, while *Aub* transcript levels were mildly increased upon H2Av GLKD, fluorescent imaging using a GFP-Aub reporter confirmed that Aubergine protein was ultimately absent in both KDs (Fig. 5C). Given the morphological defects in KD ovaries, we verified that this apparent loss of piRNA gene expression was not merely an artifact of germ cell loss. Analyzing the expression of genes known to be specifically expressed in either nurse or follicular cells showed that while somatic genes increased in relative abundance, the majority of broad germline markers remained stable (Fig. S5C). This argues against tissue distortion as the primary cause of piRNA factor loss. Furthermore, the stable expression of *vasa* and the accumulation of residual Ago3 in perinuclear foci (Fig. 5C, S5C) suggest that the nuage structure itself remains at least partially intact. Thus, piRNA factors are disproportionately and specifically dependent on H2Av and Arp6 for their expression.

To determine whether H2Av regulates these piRNA genes at the transcriptional level, we performed ChIP-qPCR using an antibody against initiating Pol II (Pol II pSer5). These assays indicated a significant loss of Pol II over several piRNA genes upon H2Av or Arp6 GLKD, supporting a mechanism of transcriptional failure (Fig. S5D) Interestingly, Pol II was also reduced at the *rhino* promoter, despite only moderate changes in its steady-state transcript levels. It is worth noting that Pol II occupancy also decreased at our control gene, *rp49*, upon H2Av KD. While this complicates baseline normalization for qPCR, it aligns with previous reports detailing H2Av’s widespread role in maintaining transcriptionally active chromatin landscapes (Ibarra-Morales et al. 2021, Hsiao et al. 2023, Chen et al. 2025). —a global promoter-level defect further corroborated by our genome-wide Pol II ChIP–seq data (explored below in Fig. 6A).

**Figure 6.**
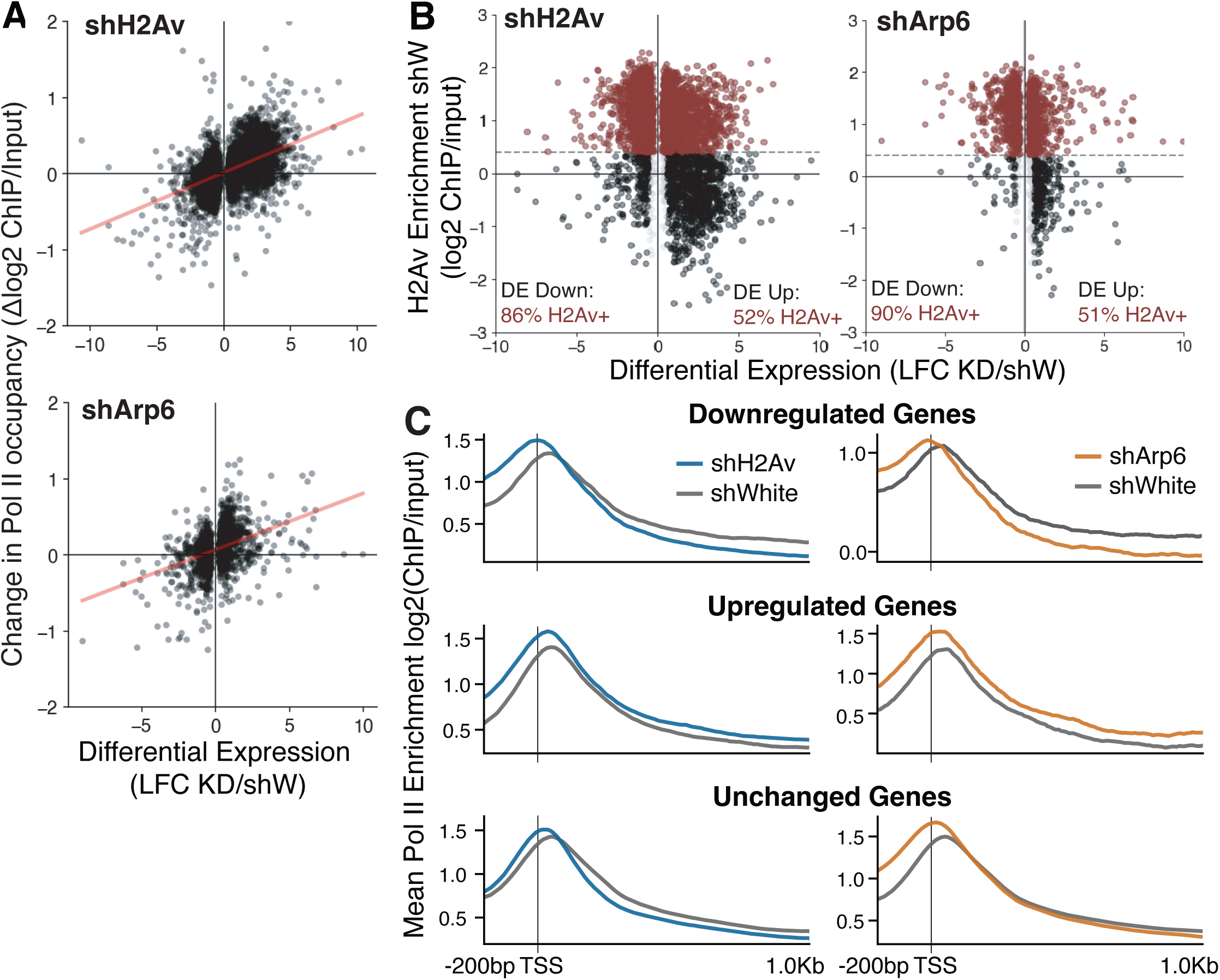
H2Av and Pol II signal at gene transcription start site in genes upregulated, downregulated, or unchanged in shH2Av knockdown. (A) Change in Poll II pSer5 ChIP signal at gene promoters correlates with change in transcript level upon H2Av (top) and Arp6 (bottom) GLKD. Each dot represents a gene, with linear regression fit shown as a red line. (B) H2Av accumulation at promoters predominantly leads to transcriptional activation. H2Av ChIP-seq signal enrichment at all significantly differentially expressed genes upon H2Av (left) and Arp6 (right) GLKD. Genes with H2Av signal around the TSS (LFC H2Av ChIP/Input > 0.4 at TSS +/- 1kb) are shown in red. % indicate genes with H2Av enrichment at TSS from the down or upregulated genes, respectively. (C) Loss of H2Av or Arp 6 alters Pol II pSer5 distribution around the TSS at genes with promoter H2Av-enrichment. Pol II pSer5 ChIP-seq signal in each expression category for respective knockdown conditions, around the gene TSS, shown as meta-profile overlay of shW (gray) with shH2Av (left, blue) and shArp6 (right, orange).

Finally, to link the transcriptional defects in the differentially expressed piRNA pathway genes directly to local H2Av deposition, we utilized H2Av ChIP-seq data from control and shArp6 ovaries. In the Arp6 KD we observed little overall change in global H2Av occupancy and no genome-wide correlation with differential expression, likely due to the moderate knockdown efficiency. However, three of the core piRNA genes downregulated in both knockdowns (*moon*, *BoYb*, and *AGO3*) exhibited a significant local decrease in promoter H2Av enrichment upon Arp6 depletion (Fig. S5E-F). We also found a loss of H2Av occupancy at the *rhino* promoter in shArp6, despite the lack of a strong corresponding transcript change in that specific knockdown (Fig. S5E). These findings supporting an H2Av-mediated effect of Arp6 on piRNA pathway gene expression. While targeted, these findings support a mechanism where Arp6 mediates the local deposition of H2Av at critical piRNA promoters to license their transcription.

### H2Av directly regulates gene promoters in the ovary, acting predominantly as an activating signal facilitating pause-release

Both H2Av and Arp6 have been implicated in the transcriptional regulation of host genes in other cellular contexts, exhibiting both activating and repressive effects depending on interacting partners (Swaminathan et al. 2005, Kotova et al. 2011, Scacchetti et al. 2020, Ibarra-Morales et al. 2021, Hsiao et al. 2023, Chen et al. 2025). This aligns with the widespread up- and downregulation of host genes we observed upon H2Av and Arp6 GLKD (Fig. 5A-B). To confirm that these expression changes were driven by transcriptional mechanisms, we compared the RNA-seq data with the Pol II pSer5 ChIP-seq data. Indeed, we found a highly significant association between promoter Pol II occupancy and global gene expression changes in the GLKDs (Fig. 6A) (Spearman’s ρ = 0.44, p = 5.1 x 10^-225^), with 63.6% of genes changing in the same direction (binomial p= 3.6 x 10^-80^). Furthermore, we observed a distinct co-occurrence of H2Av and Pol II at promoters in wild-type (shW) ovaries: 72.5% of genes with H2Av occupancy also display Pol II signal, compared to only 22% of genes lacking H2Av (Fig. S6A-B). This association was particularly pronounced among expressed genes, with 75.8% of expressed genes (mean DESeq2-normalized expression >10) falling within the quadrant of H2Av and Pol II promoter enrichment (Fisher’s exact test; odds ratio= 13.3, p <1×10^-300^), compared to only 19% of low/non-expressed genes having both H2Av and Pol II enrichment (Fig. S6B).

To definitively identify which of these differentially expressed genes are direct targets of H2Av, we performed genome-wide H2Av ChIP-seq analysis in control flies and analyzed occupancy over the TSS of all 13,986 annotated protein-coding genes. In control ovaries, ∼8,000 genes exhibited promoter-proximal H2Av accumulation (>0.4 LFC ChIP/input), representing putative direct targets. Cross-referencing these targets with our RNA-seq data revealed a striking pattern: while ∼52% of upregulated genes were H2Av-enriched (matching the ∼58% genome-wide average), an overwhelming 86–90% of genes downregulated in either shArp6 or shH2Av ovaries exhibited promoter H2Av occupancy (Fig. 6B). Consistent with this trend, downregulation was ∼4–7-fold more frequent among H2Av-enriched genes than non-enriched genes, whereas upregulation showed no such bias. This indicates that promoter-associated H2Av directly regulates its genomic targets in the ovary, acting predominantly as a positive driver of transcription.

Notably, the topological positioning of H2Av did not dictate a gene’s expression class. Across all groups, H2Av displayed a similar bimodal distribution around the TSS, consistent with a canonical nucleosome-free region (NFR), with no significant differences in peak position or shape (Fig. S6C-D). Instead, genes dependent on H2Av for expression were distinguished primarily by their signal intensity. Among H2Av-enriched targets, baseline promoter H2Av levels differed significantly across expression classes (Kruskal–Wallis H = 62.3, p = 3.0 × 10⁻¹⁴). Downregulated genes showed the highest H2Av enrichment, unchanged genes had intermediate levels, and upregulated genes had the lowest (all BH-corrected q < 10⁻⁵) (Fig. S6E). Thus, H2Av acts primarily as an activating signal, and its requirement for robust transcription scales directly with its local concentration at the promoter. Repressive functions of H2Av appear to be either secondary, indirect, or restricted to loci where the H2Av signal is relatively weak.

Because H2Av has previously been implicated in regulating promoter-proximal pausing and facilitating the transition of Pol II into the elongation phase (Weber et al. 2014), we asked if the observed transcriptional collapse could be explained by a defect in Pol II dynamics. We characterized the Pol II pSer5 signal distribution specifically at the promoters of H2Av-enriched genes, comparing the knockdowns to the shW control using basic metrics of peak position, width, and height.

In control ovaries, this metagene analysis revealed a canonical promoter-proximal Pol II peak centered ∼20 nt downstream of the TSS (μ ≈ +17.5 bp), consistent with normal Pol II pausing upstream of the +1 nucleosome prior to elongation (Kwak et al. 2013, Jonkers and Lis 2015, Day et al. 2016). However, in both knockdowns, the Pol II peak at downregulated genes underwent a structural distortion (Fig. 6C). The peak narrowed significantly (∼10 bp width reduction, p < 10⁻¹²) and shifted sharply upstream by 16.7 bp in the Arp6 KD (p = 1.27 × 10⁻^4^) and 17 bp in the H2Av KD (p = 1.65 × 10⁻¹²) (Table 1.). Critically, this upstream shift repositions the Pol II peak directly over the TSS (∼0 bp), indicating that while Pol II is successfully recruited to the initiation site, it stalls and fails to reach the standard pause-release checkpoint. Unchanged genes exhibited a similar, albeit much weaker, upstream shift and narrowing.

**Table 1.** Quantification of Pol II pSer5 signal distribution across the TSS of H2Av-positive genes, categorized by change in expression upon H2Av or Arp6 GLKD. The change in the center-of-mass (COM) position and width, and the height of the peak are calculated for each gene then the median and p-value were calculated across each expression category.

In contrast, upregulated genes displayed an entirely different Pol II profile. While the Arp6 GLKD showed no significant change in peak position or width for these genes, the H2Av GLKD caused a 17-bp *downstream* shift (p = 4.24 × 10⁻¹⁰), moving the peak deeper into the gene body (∼+34 bp). This implies an uninhibited release from pausing in the absence of H2Av at these specific loci (Fig. 6C). Finally, across all classes of genes and in both knockdown conditions, the Pol II peak significantly increased in absolute height relative to shWhite (+0.048–0.16, all highly significant) (Table 1.).

Overall, these structural changes in Pol II occupancy perfectly mirror the observed changes in gene expression. The increased peak height combined with a profound upstream shift at downregulated targets indicates that Pol II piles up and stalls exactly at the TSS. Conversely, the downstream shift at upregulated genes suggests premature or uninhibited pause release at genes where transition into active elongation is normally repressed by H2Av. Thus, H2Av serves as a critical rheostat for Pol II pause dynamics.

## Discussion

### H2Av and Arp6 silence TEs in the ovarian germline primarily through the piRNA pathway

Unlike mammalian systems where the evolutionary paralogs H2A.Z and H2A.X divide labor between transcriptional regulation and the DNA damage response, respectively, *Drosophila* H2Av coordinates both physiological processes simultaneously (Baldi and Becker 2013, Scacchetti and Becker 2021), making experimental separation of these functions often challenging. Even the transcriptional function of H2Av seems complex, featuring context-dependent roles in both gene activation and silencing. For instance, *Drosophila* H2Av is strictly required for the initial establishment of constitutive heterochromatin and the maintenance of euchromatic silencing (Leach et al. 2000, Swaminathan et al. 2005). Cytological studies reveal that H2Av localizes robustly to centromeric heterochromatin, and *His2Av* mutations act genetically as Polycomb Group (PcG) suppressors of position-effect variegation (PEV) (Swaminathan et al. 2005). Mechanistically, the incorporation of H2Av licenses the histone acetyltransferase Chameau to acetylate H4K12, a prerequisite for subsequent H3K9 methylation and HP1 recruitment at centromeres and ectopic heterochromatin loci (Grienenberger et al. 2002, Swaminathan et al. 2005). *In vitro* studies further support a role for H2A variants in chromatin condensation, demonstrating that the H2A.Z acidic patch alters the nucleosome surface to facilitate intermolecular interactions and HP1 alpha binding (Fan et al. 2002, 2004). At the level of transposon control, the N-terminus of H2Av is critical for PARP recruitment and the subsequent silencing of *Copia* and *Gypsy* retroelements (Tulin et al. 2002, Kotova et al. 2011). Interestingly, H2Av nucleosomes are intrinsically positioned on the inverted repeats of DNA-TIR transposable elements (such as *1360*), potentially facilitating their transcription, whereas this organization is absent at retrotransposon-related elements (Zhang and Pugh 2011), indicating that H2Av’s role in transposon regulation may be TE-specific. Taken together, current literature shows that H2Av is a critical factor for heterochromatin formation, the silencing of heterochromatic genes, and TE regulation. Consistent with these repressive functions, previous genome-wide screens identified H2Av and Arp6 as essential factors for TE silencing, demonstrating sterility and the derepression of *HetA*, *TAHRE*, *Blood*, and *Burdock* TEs, as well as a *Gypsy* LTR-LacZ reporter (Czech et al. 2013, Handler et al. 2013). Despite these genetic links, mechanistic studies have been lacking. Our H2Av ChIP-seq analysis revealed no accumulation—and indeed, an apparent depletion—over most TE sequences. While this depletion may partially reflect the known reduced solubility of highly compacted heterochromatin during standard sonication (Becker et al. 2017), the overall lack of H2Av enrichment is consistent with the relatively moderate Pol II increase we observe for most TE families upon GLKD. This argues against strong, direct regulation by H2Av at the TE loci themselves (Fig. 2A).

Instead, our data reveal that H2Av and Arp6 suppress TEs indirectly by maintaining the piRNA pathway. We observed a severe, coordinated downregulation of genes spanning the entire piRNA biogenesis and effector cascade. This includes factors required for precursor transcription (*cutoff* and *Trf2*, which is recruited by the RDC complex to license transcription); cytoplasmic biogenesis (the Piwi protein *Ago3* and the mitochondrial scaffolding protein *Gasz*); and nuclear effector functions (the E3 SUMO ligase *Su(var)2-10*). *Maelstrom*, which is implicated across all these processes, was one of the most strongly affected genes. Notably, the loss of Rhi protein was far more severe than its mRNA reduction, likely due to the destabilization of Rhi upon the loss of its binding partner, Cuff (Mohn et al. 2014, Chen et al. 2021).

As a result of this transcriptional collapse, we observe a global loss of mature piRNAs (Fig. 2C) due to the depletion of RDC from dual-strand clusters, the collapse of piRNA biogenesis from the major *42AB* cluster, and a slightly reduced ping-pong signature upon H2Av GLKD. Thus, the global TE derepression we observe is driven by this profound loss of piRNAs, evidenced by the strict correlation between derepressed TEs and the loss of their cognate targeting piRNAs (Fig. 2D-E). Importantly, while cluster transcription is severely impaired, we observed neither H2Av enrichment (Fig. S3E), nor a loss of H3K9me3 at these loci. This supports an indirect regulatory mechanism and argues against H2Av directing H3K9me3 deposition at clusters, as has been proposed for centromeric regions (Swaminathan et al. 2005). In contrast, Rhi levels were strongly reduced across all dual-strand clusters, and nuclear Cuff and Rhi foci vanished entirely (Fig. 4A-B, Fig. S4A-C). Notably, while the reduction of *rhino* mRNA was only significant only upon H2Av GLKD (Fig. 5B), we do observe a substantial reduction in Pol II occupancy over the *rhino* gene (Fig. S5E) alongside diminished Rhi protein levels in both GLKDs (Fig. S4E).

How piRNA pathway genes are disproportionately reliant on H2Av-mediated activation remains an open question. This enrichment is not an artifact of a global decrease in germline-specific genes caused by morphological changes, as other broad germline markers (*vasa*, *oskar*, *bam*) remained stable (Fig. S5C). Previous studies demonstrate that the expression of piRNA pathway genes can be orchestrated by specific transcription factors, such as A-myb in mouse and rooster testes (Li et al. 2013, Yu et al. 2023), or Ovo in *Drosophila* and vertebrate ovaries (Alizada et al. 2025). It is possible that H2Av, or a co-factor enhancing its activating role, is recruited to piRNA gene promoters by such sequence-specific TFs. Alternatively, H2Av may regulate a master-regulator TF that in turn coordinates the expression of multiple piRNA genes. While motif searches did not identify an enrichment for Ovo binding sites (Alizada et al. 2025), we did identify binding sites for Caudal, Elba2, and DREF (a factor previously implicated in regulating piRNA expression; (Kobelyatskaya et al. 2025) in the promoters of our most significantly downregulated piRNA genes. Furthermore, all three TFs are expressed in the fly ovary, and the expression of *Caudal* and *Elba2* is significantly reduced upon GLKD of both H2Av and Arp6. This raises the exciting possibility that these factors could be master regulators orchestrating piRNA gene expression in the *Drosophila* female germline, a hypothesis that warrants future exploration.

### H2Av regulates telomeric transposons independently of its piRNA effects

The striking exception to the indirect, piRNA-mediated TE regulation by H2Av and Arp6 occurs at the telomeric TEs: *HetA*, *TART*, and *TAHRE* (*HTTs*). In contrast to most TEs, *HTTs* exhibited distinct H2Av enrichment (Fig. 2A), consistent with classical cytological observations of robust H2Av banding at polytene chromosome telomeres (Leach et al. 2000, Rong 2008). Upon H2Av depletion, *HTTs* suffered a severe loss of the repressive H3K9me3 mark and were moderately but significantly derepressed, despite exhibiting little-to-no discernible loss of their targeting piRNAs (Fig. S2C). These findings point to a direct, chromatin-based regulatory mechanism at chromosome ends. How H2Av is specifically recruited to telomeric TEs and why it regulates them through a mechanism distinct from other transposons, remain open questions.

Previous studies established that the piRNA pathway works in tandem with local chromatin factors to safeguard telomeres, and that impairment of the piRNA pathway alone is sufficient to derepress *HTTs* (Savitsky et al. 2006, Klenov et al. 2007, Shpiz et al. 2007, 2011, Lim et al. 2009, Radion et al. 2018, Teo et al. 2018, Iyer et al. 2023). However, our data suggest that H2Av operates upstream of, or parallel to, these piRNA-dependent effects at the telomere. Furthermore, our RNA-seq data revealed that H2Av depletion downregulates host genes critical for telomere maintenance. For instance, we observed a loss of *polo* kinase, which is required to maintain transcriptional silencing at telomeric boundaries (Morgunova et al. 2021). Thus, H2Av appears to safeguard telomeric integrity through a diverse, multi-layered mechanism involving direct nucleosomal incorporation, the maintenance of H3K9me3, and the transcriptional support of protective host factors.

### H2Av has a complex effect on host gene regulation

Moving beyond TEs to the host genome, our data underscore H2Av’s primary role as a positive transcriptional regulator. In budding yeast, the H2A.Z paralog occupies both the -1 and +1 nucleosomes flanking the nucleosome-free region (NFR), directly regulating start site selection (Guillemette et al. 2005, Raisner et al. 2005, Zhang et al. 2005, Raisner and Madhani 2006, Mavrich et al. 2008a). Previous studies suggested that *Drosophila* generally lacks H2Av at the -1 nucleosome, and the entire genic array of nucleosomes is shifted downstream by ∼75 base pairs (Mavrich et al. 2008b), allowing unimpeded assembly of the Pol II initiation complex within the NFR. While our bulk ChIP-seq data lacks single-nucleotide resolution to pinpoint exact nucleosome positioning, the H2Av localization we observe around ovarian promoters is consistent with the +1 nucleosome residing downstream of the TSS, enabling Pol II access. However, our high-throughput data also reveals a strong enrichment of H2Av *upstream* of the NFR that is comparable to its downstream enrichment. This implies that, similar to yeast, *Drosophila* ovaries incorporate H2Av into both the +1 and -1 nucleosomes.

Consistent with the dual activating-repressing function of H2Av (Swaminathan et al. 2005, Kotova et al. 2011, Ibarra-Morales et al. 2021, Hsiao et al. 2023, Chen et al. 2025), our genome-wide analysis revealed widespread changes in host gene expression upon GLKD of either H2Av or Arp6. Strikingly, 86–90% of the genes downregulated in KD ovaries possessed H2Av at their promoters, indicating they are putative direct targets. These results align with previous findings that H2Av incorporation into promoter-proximal nucleosomes is required for transcription (Mavrich et al. 2008b, Weber et al. 2014); for instance, a recent study demonstrated that the Tip60-H2Av axis is strictly required for Notch signaling activation during *Drosophila* development (Chen et al. 2025). Importantly, we observed a quantitative difference in local H2Av levels: baseline H2Av signal differed significantly across expression classes, with the genes most dependent on H2Av for their expression exhibiting the highest wild-type enrichment. Supported by the distinct co-occurrence of H2Av, active Pol II, and high transcript expression in control ovaries, our analyses indicate that H2Av’s direct role is primarily as an activating signal, whereas its repressive functions are either secondary or restricted to loci where the promoter H2Av signal is relatively weak.

We note that based on bulk ChIP-seq data, it is impossible to differentiate whether the higher overall H2Av signal at these activated genes represents an increased density of homotypic (H2Av/H2Av) nucleosomes, a greater total number of H2Av-bearing nucleosomes, or simply a higher fraction of cells with H2Av-bearing promoters. Functionally, these scenarios could yield significantly different outcomes. Previous mapping demonstrated that homotypic H2Av nucleosomes resemble classical active chromatin and are enriched downstream of active promoters, whereas they are depleted downstream of paused polymerases (Weber et al. 2010). Conversely, PARP1 activation—which creates chromatin permissive to Pol II loading—requires the interaction of neighboring nucleosomes, where one contains canonical H2A and the other variant H2Av (Kotova et al. 2011). In such cases, relatively lower levels of H2Av are sufficient to ensure the presence of these heterotypic contexts. Finally, the H2A variant has been proposed to keep chromatin poised for activation by promoting local nucleosome folding while preventing dense, large-scale compaction (Fan et al. 2002). In a distinct boundary role, H2Av protects active euchromatic genes by physically blocking the spread of silencing from neighboring heterochromatin (Meneghini et al. 2003). Thus, the quantitative differences in H2Av occupancy we observe possibly represent diverse, context-dependent regulatory mechanisms through which H2Av governs gene expression.

### H2Av dictates Pol II promoter escape and pause-release dynamics

How exactly does H2Av regulate transcription at H2Av-dependent and H2Av-silenced genes? In agreement with previous findings (Mavrich et al. 2008b), we observed that at active genes in control ovaries, Pol II pSer5 accumulates in a paused state centered ∼20 bp downstream of the TSS, immediately upstream of the H2Av signal at the +1 nucleosome (Fig. S6B). However, upon H2Av or Arp6 depletion, the Pol II landscape at H2Av-dependent (downregulated) genes is severely distorted. The Pol II peak narrows, increases in amplitude, and shifts ∼17–23 bp upstream, stalling exactly at the transcription initiation site (∼0 bp). Concurrently, the Pol II signal is substantially reduced downstream across the gene body (Fig. 6C).

This upstream shift at the TSS coupled with the depletion of elongating Pol II implies that H2Av might facilitate both promoter escape and the subsequent release from pausing. Canonical H2A nucleosomes form a rigid barrier to the transcription machinery (Weber et al. 2014). The incorporation of H2Av into the +1 nucleosome counteracts this by promoting nucleosome "unwrapping," thereby lowering the kinetic barrier for the passing polymerase (Wen et al. 2020, Li et al. 2023). Depletion of H2Av decreases this nucleosomal plasticity at the promoter, causing Pol II accumulation—a mechanical bottleneck consistent with the increased Pol II ChIP signal amplitude we observed at downregulated genes (Fig. 6C).

Interestingly, the distinct redistribution of the Pol II signal from the +1 nucleosome back to the TSS upon GLKD suggests two non-mutually exclusive mechanisms. First, the narrowed, upstream-shifted Pol II peak could reflect the backtracking of the polymerase upon encountering an impassable nucleosomal barrier (Weber et al. 2014). Alternatively, it may signify a profound defect in transcription initiation and promoter clearance, as observed at the inducible *Drosophila hsp70* gene. At the hsp70 gene promoter, in the absence of the H2Av-exchange complex dTip60 (which contains Arp6), Pol II can successfully assemble, but fails to release upon induction, stalling directly at the TSS (Kusch et al. 2014). This initiation-clearance defect provides a compelling explanation for the increased Pol II ChIP amplitude at our downregulated targets: the inability to escape the TSS forces newly recruited polymerases to continuously pile up without successfully transitioning into elongation, yielding a taller, narrower ChIP peak coupled with a severe loss of mature transcripts.

## Materials and methods

### Fly stocks

Flies were maintained at 25°C. For ovary dissections, females were aged for 1-3 days and fed yeast paste for 24 hours prior to dissection. All fly lines containing the UASp promoter sequence were driven by *nos-GAL4-VP16*.

Flies carrying an shRNA targeting *His2Av* (BDSC #34844) and the *nos-GAL4-VP16* driver (BDSC #4937) were obtained from the Bloomington *Drosophila* Stock Center. GFP-Rhi (VDRC #313340), GFP-Cuff (VDRC #313269), and *w[1118]* (VDRC #60000) lines were obtained from the Vienna *Drosophila* Resource Center. Transgenic flies expressing GFP-Aub were generated as described previously (Webster et al., 2015). The mKate2-AGO3 knock-in fly was a kind gift from Toshie Kai (Lin et al. 2023). Flies harboring *nos-Gal4-VP16* on the second chromosome were a gift from Trudi Schüpbach (Pane et al. 2011). GFP-Mael was a gift from Gregory Hannon.

### Generation of transgenic flies

All transgenic flies were generated via phiC31-mediated recombination (BestGene Inc.). To generate *Arp6*-KD flies, a short hairpin (Table S1.) containing sequences against *Arp6* was ligated into the pValium20 vector (Ni et al. 2011) (*Drosophila* Genomics Resource Center (DGRC) Stock 1467; https://dgrc.bio.indiana.edu//stock/1467; RRID:DGRC 1467). The resulting construct was integrated into the attP2 landing site (genotype: *y[1] w[67c23]; P{CaryP}attP2*, BDSC #8622). To obtain the *white*-KD control line, a hairpin (Table S1.) targeting the *white* gene was ligated into the pValium22 vector (Ni et al., 2011) (DGRC Stock 1468; https://dgrc.bio.indiana.edu//stock/1468; RRID:DGRC 1468) and integrated into the attP40 landing site (genotype: *y[1] w[67c23]; P{CaryP}attP40*).

To generate the mCherry-Rhi construct under the control of its endogenous promoter, mCherry DNA and a ∼2 kb region upstream of the *rhi* gene (containing the putative *rhi* promoter) were PCR amplified, introducing KpnI/AgeI and StuI/KpnI restriction sites, respectively. The two PCR fragments were digested and ligated into a modified pPGW Gateway destination vector (pUASp-eGFP-GW-phiC31), wherein the pUASp promoter and eGFP sequences were replaced with the *rhi*-promoter and mCherry sequences. Rhi cDNA was then inserted into the resulting destination vector (pRhi-mCherry-GW-phiC31) via TOPO cloning (Thermo Fisher Scientific, Cat# K240020) and a Gateway LR clonase reaction (Thermo Fisher Scientific, Cat# 11791020). The final construct was integrated into the attP40 landing site. To generate the Arp6-GFP construct under the control of the pUASp promoter, Arp6 cDNA was PCR amplified (Suppl. Table 1), cloned into the pENTR™/D-TOPO™ vector, and subsequently moved into a modified pPWG Gateway destination vector (pUASp-GW-eGFP-phiC31) containing a mini-*white* marker via a Gateway LR clonase reaction. This construct was also integrated into the attP40 landing site.

### Fertility test

To test female fertility for each genotype (*white*-KD, *Arp6*-KD, and *His2Av*-KD), single 1-day-old virgin females were mated with two 3-day-old *w[1118]* males for 5 days (n=10 parallel crosses per genotype). To test male fertility, single 3-day-old knockdown males were mated with one 2-day-old *w[1118]* virgin female for 5 days (n=10 parallel crosses per genotype). Adults were discarded from the vials after 5 days, and all eclosed progeny were counted 14 days later (19 days post-mating). Two independent biological replicates were performed (n=2).

### RNA *in situ* Hybridization Chain Reaction (HCR)

For *rhi* (unique identifier: 3682/D707) and *cuff* (unique identifier: 5486/G063) RNA *in situ* hybridization chain reactions, probes, amplifiers, and buffers were purchased from Molecular Instruments (molecularinstruments.org). RNA *in situ* HCR v3.0 (Choi et al. 2018) was performed according to the manufacturer’s instructions for generic samples in solution. Following hybridization, ovaries were washed, counterstained with DAPI, mounted, and imaged via confocal microscopy as described below.

### Confocal microscopy and image processing

Ovaries from knockdown flies expressing GFP-Cuff, GFP-Rhi, mKate2-AGO3, GFP-Aub, and GFP-Mael were dissected in ice-cold phosphate-buffered saline (PBS) and fixed in PBST (PBS containing 0.1% Tween-20) supplemented with 4% formaldehyde for 20 minutes at room temperature (RT). Ovaries were then washed three times in PBST at RT, with DAPI (1:1000) added during the second wash. Samples were mounted in VECTASHIELD Antifade Mounting Medium (Vector Laboratories, Inc., Cat# H-1000-10). Images, including samples from RNA *in situ* HCR, were acquired using a Leica Stellaris 8 FALCON confocal microscope with a 63x oil immersion objective. Identical acquisition parameters were used for control and GLKD ovaries, and images were processed using ImageJ (Schneider et al. 2012).

### RNA extraction and RT-qPCR

Ovaries from control (shWhite) and *Arp6* or *H2Av* germline knockdown flies expressing GFP-Cuff, GFP-Rhi, mKate2-AGO3, GFP-Aub, and GFP-Mael were dissected in ice-cold PBS. Total RNA was extracted from 10-20 pairs of ovaries using TRIzol Reagent (Invitrogen, Cat# 15596026). Extracted total RNA (1 µg) was treated with DNase I (Invitrogen, Cat# 18068-015), and first-strand cDNA synthesis was performed using random hexamers and SuperScript III Reverse Transcriptase (Invitrogen, Cat# 18080-044). qPCR was performed on a Mastercycler ep realplex 2 system (Eppendorf) using MyTaq HS Polymerase Mix (Thomas Scientific, Cat# BIO-25046) and SYBR Green I stain (Invitrogen, Cat# S7563) according to the manufacturers’ instructions. Target gene expression was normalized to the geometric mean expression of two reference genes, *rp49* and *GAPDH1*. Experiments were conducted using at least two biological replicates and three technical replicates. Primers used for qPCR are listed in Table S1.

### RNA-seq

Total RNA was extracted from dissected ovaries using TRIzol Reagent. Approximately 1.5 µg of total RNA was depleted of ribosomal RNA using the *Drosophila melanogaster* riboPOOL Kit (Galen Molecular, Cat# dp-K006-00007). RNA-seq libraries were subsequently prepared using the NEBNext Ultra II Directional RNA Library Prep Kit for Illumina (NEB, Cat# E7765S) according to the manufacturer’s instructions. Libraries were sequenced on an Illumina NextSeq 2000 instrument. Two biological replicates were sequenced for all genotypes.

### Small RNA-seq

Small RNA (sRNA) was extracted from 15-20 pairs of ovaries using the TraPR Small RNA Isolation Kit (Lexogen). sRNA-seq libraries were prepared using the NEBNext Small RNA Library Prep Set for Illumina (Multiplex Compatible) (NEB, Cat# E7330S) according to the manufacturer’s instructions. Libraries were sequenced on an Illumina NextSeq 2000 instrument. Two biological replicates were sequenced for all genotypes.

### ChIP-qPCR and ChIP-seq

Chromatin immunoprecipitation (ChIP) experiments were carried out as previously described (Le Thomas et al. 2014). Pre-cleared chromatin samples were immunoprecipitated overnight at 4°C using anti-His2Av (Active Motif, Cat# 61751), anti-Pol II (phospho-Ser5; abcam, Cat# ab5408), anti-GFP (Invitrogen, Cat# A-11122), or anti-H3K9me3 (abcam, Cat# ab8898) antibodies. Immunoprecipitated complexes were incubated with 50 µl of Dynabeads Protein G for 5h at 4°C. Beads were then washed five times for 10 min each in LiCl wash buffer (10 mM Tris pH 7.4, 500 mM LiCl, 1% NP-40, 1% sodium deoxycholate). Samples were subsequently incubated with 100 μg of proteinase K in PK Buffer (200 mM Tris pH 7.4, 25 mM EDTA, 300 mM NaCl, 2% SDS) at 55°C for 2 h, followed by 65°C overnight. DNA was purified via phenol-chloroform extraction.

ChIP-qPCR was performed on two biological replicates for all genotypes using a Mastercycler ep realplex 2 system (Eppendorf) with MyTaq HS Polymerase Mix and SYBR Green I stain. All ChIP-qPCR signals were normalized to their respective inputs. Primers used for ChIP-qPCR are listed in Table S1. ChIP-seq libraries were prepared using the NEBNext Ultra II DNA Library Prep Kit for Illumina (NEB, Cat# E7103S) according to the manufacturer’s instructions. Two biological replicates per genotype were sequenced on an Illumina NextSeq 2000 instrument.

#### Bioinformatic Analysis

##### Total mRNA-seq

Total rRNA-depleted RNA-seq reads were adapter-trimmed and quality-filtered using *Cutadapt* (v2.6). *FastQC* (v0.12.1) was used to verify expected size distributions, adapter content, and overrepresented sequences.

###### Gene expression

Total RNA-seq reads were adapter-trimmed and quality-filtered using *Cutadapt (v2.6). FastQC (v0.12.1)* was used to verify expected size distributions, adapter content, and overrepresented sequences. Filtered reads were quantified against the *Drosophila melanogaster* (dm6) transcriptome using the *Salmon* pseudo-aligner *(v1.10.3)* with default parameters. Normalization and differential expression analyses were performed using *DESeq2 (v1.42.1)*. Statistical correlation in differential gene expression between the two knockdown conditions was assessed using Spearman’s rank correlation coefficient (ρ) with a two-sided significance test. Genes with an adjusted p-value (padj) < 0.05 and |log₂FC| > 0.58 were classified as differentially expressed for downstream analyses.

To determine if the 51 established piRNA pathway genes were significantly enriched among downregulated targets, we performed a permutation test (10,000 permutations). The observed number of downregulated piRNA factors was compared to a null distribution generated by randomly sampling 51-gene sets from the entire pool of genes analyzed in DESeq2. The empirical p-value was calculated as the fraction of permutations yielding a downregulated count greater than or equal to the observed value.

###### TE expression

Trimmed RNA-seq reads were mapped to the dm6 genome using *Bowtie2 (v2.5.4)* with a high multi-mapper tolerance (-k 100) to retain reads mapping to repetitive elements. Alignments were sorted and indexed using *SAMtools*. Transcript-level TE quantification was performed using *TEtranscripts*, an algorithm optimized for multi-mapped alignments and repetitive genomic regions, utilizing the dm6 RepeatMasker annotations. Differential TE expression across conditions was quantified using *DESeq2* to calculate fold changes and statistical significance. As with host genes, Spearman’s rank correlation with a two-sided significance test was used to evaluate the consistency of differential TE expression between the knockdowns.

###### piRNA Precursor expression

Based on established *Drosophila* ovary piRNA cluster annotations, we generated non-overlapping 5-kb intervals across each genomic cluster; the final two windows of a cluster were merged if the terminal window was less than 3 kb. RNA-seq reads were aligned using *Bowtie2* (v2.5.4; unique mappings only) and sorted/indexed via *SAMtools (v1.19.2).* Reads mapping to each window were quantified non-redundantly using *featureCounts (v2.0.8)* and normalized by library size for cross-sample comparison. Differential analysis and statistical evaluation of these windows were performed using *DESeq2*.

##### Small RNA-seq

###### Filtering and characterization of small RNA classes

Small RNA libraries were adapter-trimmed and quality-filtered using *Cutadapt (v2.6)*, followed by quality control with *FastQC (v0.12.1)*. Reads were aligned to the dm6 genome using Bowtie (v1.3.1) with a 3-mismatch tolerance. We subsequently filtered out alignments to publicly available annotations of rRNA, tRNA, snRNA, and snoRNA (1-mismatch tolerance). The retained reads were mapped to miRNA sequences from *miRBase* (v22; 1-mismatch tolerance) to generate miRNA counts, which served as a robust normalization factor across samples in subsequent piRNA analyses. To control for potential degradation artifacts or shRNA contamination, small RNAs were also aligned to targeted transcripts (3-mismatch tolerance). Finally, the library was size-fractionated using *Cutadapt*, retaining putative piRNAs 23–29 nt in length.

###### piRNA origin and target analysis

To investigate differential piRNA production from cluster precursors, piRNA libraries were re-mapped to the dm6 genome using *Bowtie*, retaining only unique alignments with 0 mismatches. These alignments were quantified across the previously defined 5-kb cluster windows using *featureCounts*. The raw count matrix was filtered to remove low-abundance windows prior to analysis. Differential expression was assessed at the window level using *DESeq2*, providing the total miRNA read counts as size factors to ensure consistent cross-sample normalization. These results were further validated manually using a custom script to aggregate and normalize counts based on miRNA levels.

To evaluate piRNA phasing, genome-aligned reads were used to calculate the distance from the 3’ end of a given piRNA to the 5’ end of the nearest downstream piRNA on the same strand. Phasing strength was summarized by the Z₁-score, defined as the Z-score of the 1-nt distance signal relative to a background distribution (2–50 nt).

To analyze the TE-silencing capacity of differentially expressed piRNAs, libraries were mapped antisense to the full *dm6 RepeatMasker (v4.0.6)* TE repertoire using *Bowtie* (up to 3 mismatches). Multi-mapped reads were randomly assigned to a single position (-v 3 -M 1). TE targeting was quantified using a custom script that counts antisense piRNAs per TE family, normalized by sample miRNA levels. These trends were corroborated by mapping piRNAs antisense to *Dfam (v3.9)* TE consensus sequences (up to 3 mismatches), again randomly assigning multi-mapping reads.

For ping-pong cycle analysis, piRNA reads were mapped to TE consensus sequences (*Bowtie 1*) allowing up to 3 mismatches for antisense alignments and 0 mismatches for sense alignments. Pairs with overlapping coordinates on opposite strands were identified, and the overlap length was recorded. The ping-pong signature was plotted as the fraction of overlapping piRNA pairs across lengths of 1–29 nt. The Z₁₀-score was calculated using 1–9 nt and 11–29 nt as background.

##### H2Av, Pol II, H3K9me3, and Rhino ChIP-Seq Analysis

ChIP-seq reads were trimmed and quality-filtered similarly to RNA-seq datasets. Genomic mapping and quantification were customized based on the specific characteristics of the analyzed loci (gene promoters, TEs, or piRNA clusters).

###### ChIP Analysis at Genes

Pol II and H2Av ChIP-seq reads were aligned to the dm6 genome using Bowtie2, retaining only uniquely mapped reads. Alignments were sorted and indexed with *SAMtools*. Genome browser tracks (bigWig format) were generated using *deepTools (v3.5.5) bamCoverage* with a 10-nt bin size and CPM normalization and viewed in the Integrated Genome Viewer (IGV). For quantitative analysis, log₂ enrichment tracks (ChIP vs. Input) were generated using *deepTools bigwigCompare* with a pseudocount of 1. Promoter regions were defined as ±1kb from the TSS. DeepTools computeMatrix and plotProfile were used to generate heatmaps and aggregate positional profiles of ChIP signal across promoters. For quantitative promoter-level analyses, read counts overlapping promoter regions were quantified from ChIP and input BAM files using *bedtools multicov*, and replicate-averaged ChIP/Input enrichment values were subsequently calculated for each promoter region.

To classify genes as "H2Av-positive" in the wild-type (shW) condition, we titrated the ChIP/Input log₂(ChIP/Input) threshold (0.25 to 1.5) against visual confirmation in the IGV browser and compared it with MACS2 broad peak calls. This integrated approach established a threshold of 0.4 log₂(ChIP/Input) to accurately isolate genes with true H2Av binding for downstream targeting analysis.

For Pol II metagene analysis, three summary metrics were calculated from the TSS-centered (±1kb) profiles for each gene: peak position (center of mass), peak width (standard deviation), and peak height (maximum signal). Per-gene differences between KD and control conditions were calculated, median differences reported, and statistical significance determined via a two-sided Wilcoxon signed-rank test.

###### ChIP Analysis at piRNA Clusters

To account for the non-canonical transcription of many piRNA clusters, we employed a tile-based analytical approach. A 100-mer mappability track for the dm6 genome was generated using *GenMap (-K 100 -E 1 -t 5*). The genome was divided into 5-kb non-overlapping tiles, which were intersected with the mappability track to retain tiles with an average score ≥0.8. Tiles were categorized into three mutually exclusive classes with priority given to “cluster” assignment: tiles overlapping annotated piRNA clusters (±1 kb), and remaining tiles classified as either euchromatin or heterochromatin based on established chromosomal boundaries (Baumgartner et al. 2022). However, any 5-kb heterochromatin or euchromatin tiles that overlapped TE annotations (RepeatMasker), were removed to prevent TE tiles from confounding the signal changes in these “background tiles”.

While unique alignments were utilized for IGV browser visualization, the quantitative tile-based analysis utilized multi-map inclusive alignments (*Bowtie2 -k 50*) to account for high repeat content, with fractional read assignment to each genomic tile. Per-tile counts were obtained using *featureCounts*, and library size normalized ChIP/Input enrichment values were subsequently calculated for each tile. The input-normalized ChIP signal (H2Av, Pol II, H3K9me3, Rhino) for cluster tiles was then compared against other genomic regions for two biological replicates. Because "biologically true" Rhino binding requires H3K9me3 occupancy, all quantitative Rhino analyses were strictly restricted to tiles co-occupied by H3K9me3.

###### ChIP Analysis at TEs

Due to the highly repetitive nature of TEs, genome alignments were repeated with *Bowtie2* using a high multi-mapper tolerance (-k 50). Reads were fractionally assigned to each TE copy (*dm6 RepeatMasker v4.0.6*) using *featureCounts* and aggregated at the family level. Because Pol II and H2Av exert their activity at promoters, they were quantified at the TE TSS ±1kb. Conversely, because H3K9me3 and Rhino distribute broadly across regulated elements, their signals were quantified across the entire body of each TE copy (plus a ±1kb flank) and fractionally assigned to TE families.

## Data availability

The data discussed in this publication have been deposited in NCBI’s Gene Expression Omnibus and are accessible through GEO Series accession number GSE331324.

## Acknowledgements

We thank members of the Fejes Tóth and Aravin labs for discussions and Alexei Aravin in particular for feedback on the manuscript. We are grateful to Trudi Schüpbach, Gregory Hannon, the Bloomington *Drosophila* Stock Center, and the Vienna *Drosophila* Resource Center for providing fly stocks, and to the *Drosophila* Genomics Resource Center (NIH Grant 2P40OD010949) for vectors. We thank Igor Antoshechkin (Millard and Muriel Jacobs Genetics and Genomics Laboratory, Caltech) for sequencing assistance; Giada Spigolon and Andres Collazo (Biological Imaging Facility, Caltech) for confocal imaging support; Grace Shin (Molecular Technologies, Caltech) for HCR expertise; and BestGene Inc. for *Drosophila* embryo injections. Norbert Andrási was supported by the Nemko Postdoctoral Fellowship (part of the Caltech-BBE Divisional Fellowship). KFT was supported by the National Institutes of Health (R01 GM110217).

## Author contributions

K.F.T. conceived the study and wrote the manuscript, H.M.R. commented on the manuscript. N.A., Y.L. and K.F.T. performed experiments. N.A. analyzed qPCR data and H.M.R. performed the bioinformatic analyses.

## Supplemental Figure Legends

**Figure S1.**
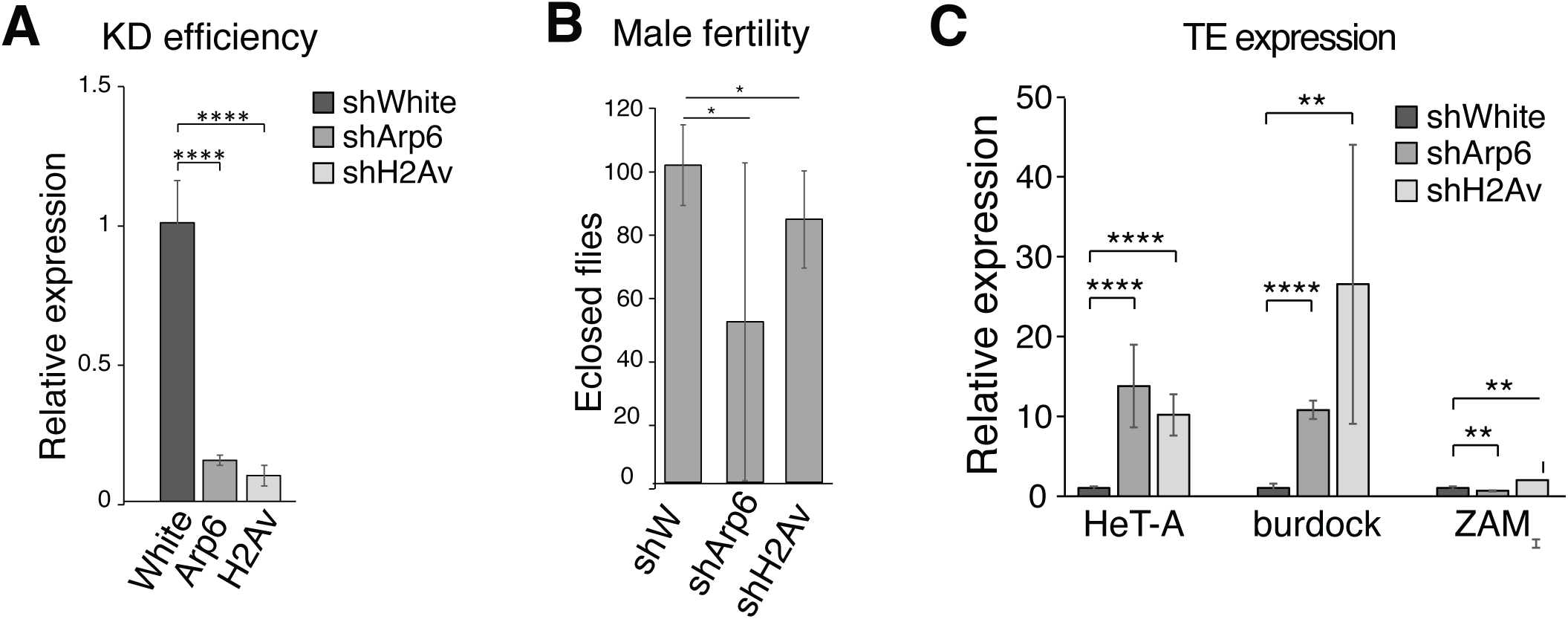
(A) RT-qPCR showing GLKD efficiency in His2Av-KD and Arp6-KD lines. Expression of His2Av and Arp6 are compared to the expression in the white-KD line. (B) Arp6 and H2Av GLKD causes slight reduction of male fertility. Respective hairpins were driven in the male germline by nos-Gal4-VP16 driver, males were crossed to wild-type females. (C) RT-qPCR analysis of select TEs indicates derepression of Het-A and Burdock upon H2Av and Arp6 GLKD. (Student’s t-test, ****p<0.0001, **p<0.01, *p<0.05; error bars represent SD)

**Figure S2.**
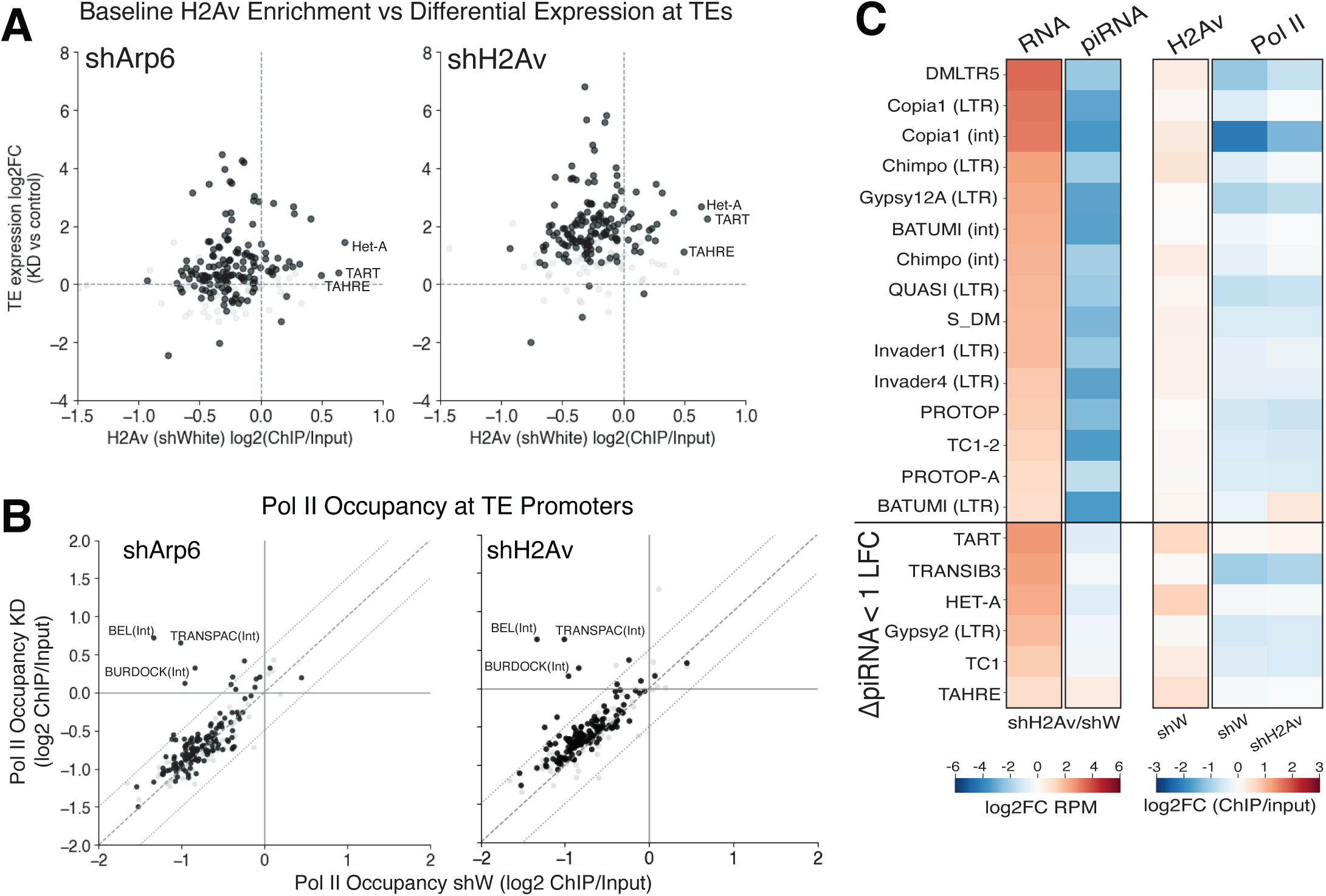
(A) TEs derepressed upon H2Av or Arp6 GLKD do not show H2Av enrichment, except for the telomeric TEs. Scatterplot showing the relationship between levels of H2Av enrichment in control ovaries (log2fc input normalized counts at TSS) vs differential TE expression upon Arp6 and H2Av GLKD. (B) Pol II pSer5 accumulation at TE promoters exhibit modest increase in a subset of TEs upon H2Av and Arp6 GLKD. Pol II pSer5 ChIP-seq signal over TE promoters in control (shW) vs H2Av (left) and Arp6 (right) GLKD. Shown is input-normalized signal average of two biological replicates. Central dashed line shows x=y, outer dashed lines show 0.5 LFC ChIP/Input. In (A) & (B) significantly derepressed TE families (padj<0.05) are shown in black (nonsignificant in grey). Significance defined based on H2Av GLKD for both conditions. (C) H2Av’s effect on TE silencing can be classified into piRNA-mediated and piRNA independent mechanisms. Shown are the 21 TEs with H2Av signal at their promoters, of which 15 show a decrease of a least 1 LFC in levels of targeting piRNA upon H2Av GLKD (top), while the other 6 do not. Fold change (log2FC shH2Av/shW) in TE transcript level based on RNA-seq (left), in piRNA level based on smallRNA-seq (middle) are indicated by differential expression color scale. Right side shows H2Av (in shW control) and Pol II pSer5 (in shW and shH2Av) ChIP-seq signal (scale showing log2 ChIP/input at the TSS +/- 1kb).

**Figure S3.**
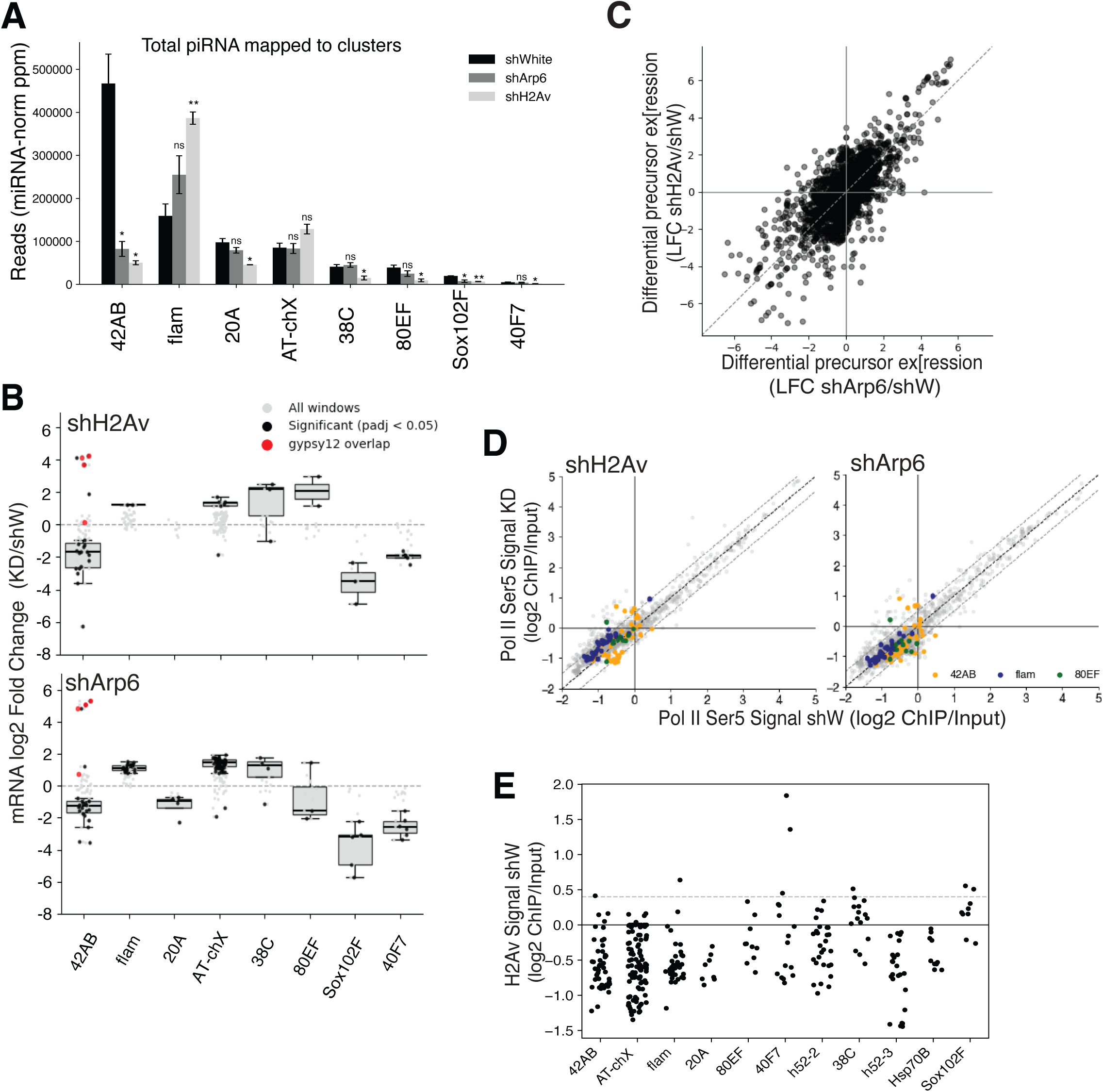
(A) piRNA abundance at select clusters in White, Arp6 and His2Av GLKD. Total piRNAs (0 mismatches, uniquely mapping) mapping to each piRNA cluster, normalized by miRNA read count for each sample. Statistical significance was determined by two-sided Student’s t-test (*p<0.05, **p<0.01; n=2 biological replicates per condition). Error bars represent SD. (B) Quantification of differential cluster precursor expression from each 5kb window in the main piRNA-producing clusters (2 replicate DESeq2 differential analysis) in shH2Av (top) and shArp6 compared to shWhite, (C) Correlation between change in piRNA cluster precursor expression in shArp6/shW vs shH2Av/shW based on the RNA-seq quantifications in 5kb windows. (D) Poll II pSer5 ChIP-seq signal enrichment over input in the H2Av GLKD (left) and Arp6 GLKD (right) compared to the White GLKD control. Each datapoint represents aggregate signal over 5kb genomic tile that overlaps any cluster annotation, averaged for 2 replicates. (E) Average H2Av ChIP-seq signal enrichment over input in the shWhite control at 5kb genomic tiles overlapping clusters.

**Figure S4.**
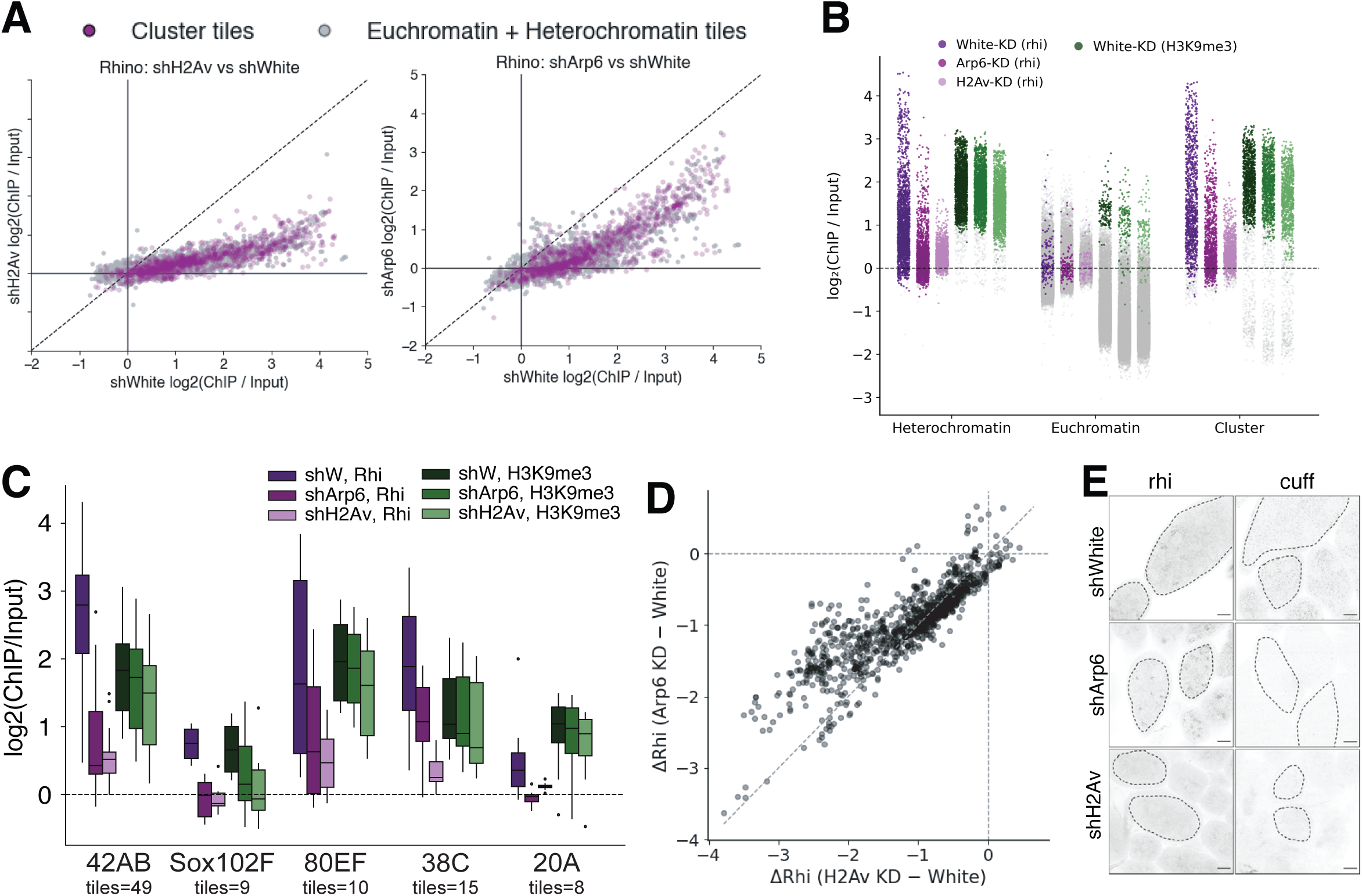
(A) H2Av (left) and Arp6 (right) GLKD lead to global loss of Rhino. Input normalized Rhino ChIP signal in genome-wide 5kb tiles. Rhino loss in tiles overlapping piRNA clusters in purple compared to background tiles in grey (i.e., non-cluster and non-TE overlapping genomic tiles). Only tiles with H3K9me3 ChIP/input signal enrichment are shown (B) Levels of input-normalized Rhino and H3K9me3 ChIP signal (2 replicate average), in clusters and remaining non-cluster, non-TE genomic tiles (split into heterochromatin and euchromatin). (C) Input-normalized H3K9me3 and Rhino ChIP enrichment at select germline clusters. (D) Correlation of the change in Rhi signal at piRNA clusters upon Arp6 and H2Av GLKD when compared to shW. Each data point represents a 5kb tile that overlaps a cluster and has H3K9me3 ChIP/input signal enrichment in shW. (E) Cuff transcript is lost upon H2Av and Arp6 GLKD, while rhi mRNA is only reduced upon H2Av depletion. Confocal images of *in situ* hybridization chain reaction signal using *rhi* and *cuff* probes. (Scale bar - 20 µm).

**Figure S5.**
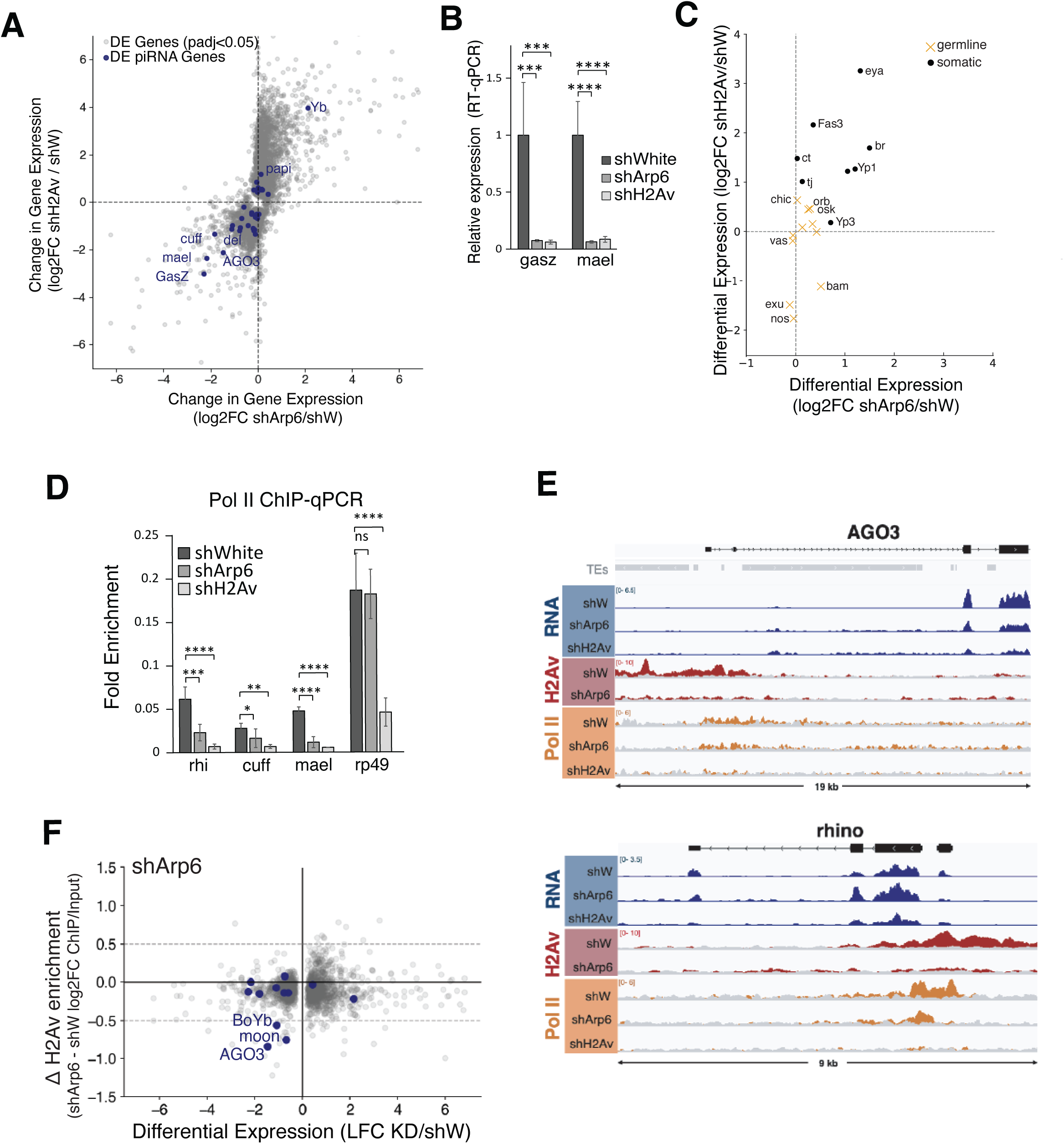
(A) Gene differential expression in shArp6 and shH2Av compared to shW is correlated. Significantly differentially expressed genes are shown in dark grey, with piRNA pathway genes highlighted in blue and labeled. (B) RT-qPCR showing loss of *gasz* and *mael* transcript in H2Av and Arp6 GLKD (Student’s t-test, ***p<0.001, ****p<0.0001, ns – not significant, error bars represent SD). (C) Differential expression (log2FC KD/shW) of select somatic (black dots) and germline (orange x) factors upon Arp6 and H2Av GLKD compared to the control. (D) Pol II Chip-qPCR of selected piRNA pathway genes and RP49, as control (Student’s t-test, *p<0.05, **p<0.01, ***p<0.001, ****p<0.0001, ns – not significant, error bars: SD). Primers were designed approx. 200bp downstream of TSS. (E) Representative single-replicate genome browser view (in IGV software) for AGO3 and Rhino showing RNA-seq expression levels in blue, H2Av ChIP-seq in red, and Pol II ChIP-seq in orange. Input signal for H2Av and Pol II ChIP are overlayed in grey. (F) Change in gene expression upon Apr6 GLKD does not correlate with H2Av loss at the TSS. Change in H2Av signal at gene TSS regions in the shArp6 knockdown plotted against the change in gene expression in shArp6. piRNA pathway genes are highlighted in blue.

**Figure S6.**
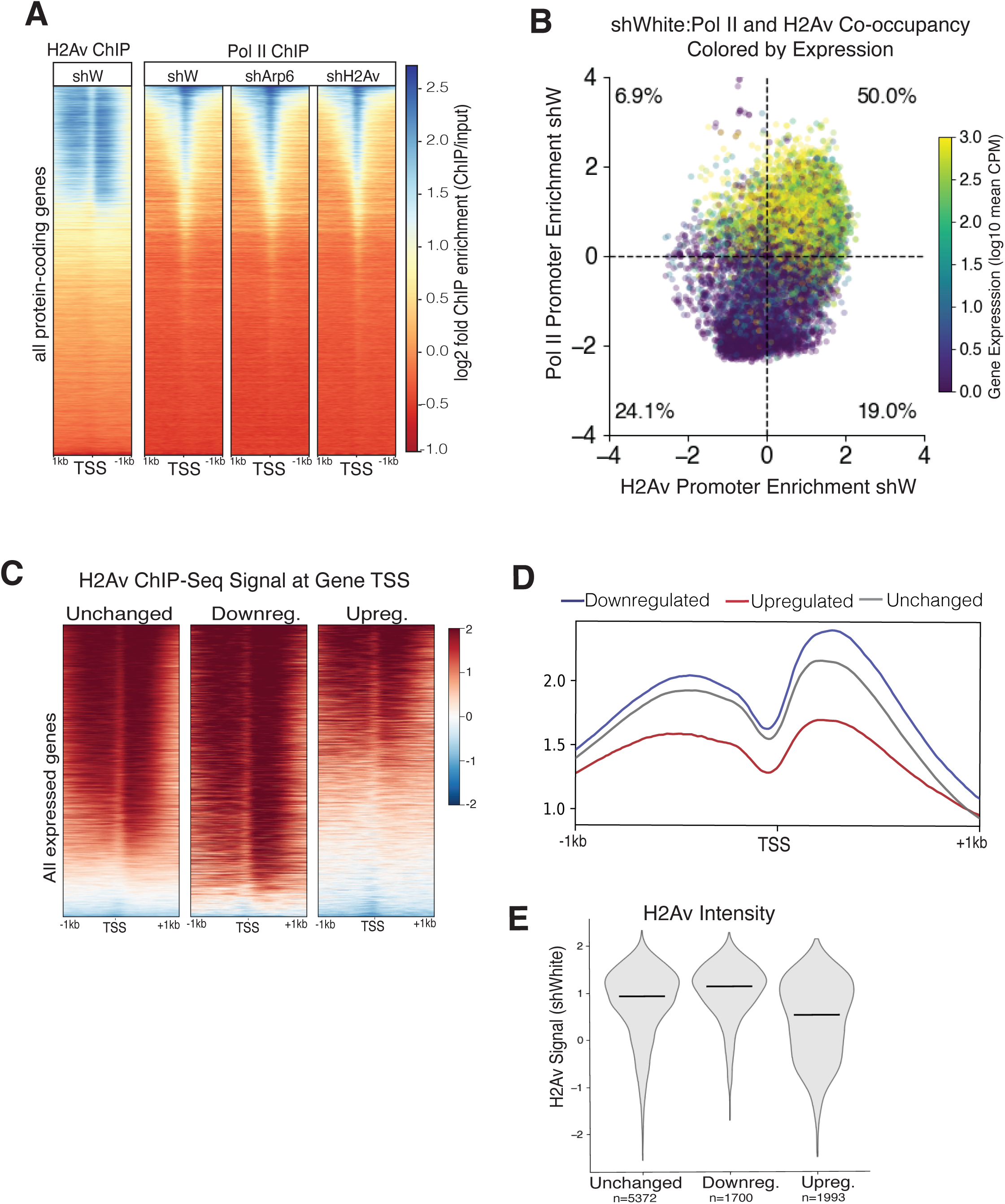
(A) H2Av and Pol II pSer5 accumulation correlate at TSSs of protein coding genes. Heatmap showing input normalized H2Av and Pol II pSer5 occupancy across all protein coding genes, averaged for 2 biological replicates. (B) Quantification of frequent co-occurrence of Pol II pSer5 and H2Av ChIP signal (input normalized) surrounding the TSS of protein coding genes in the shW control. Percents displayed reflect the quantity of genes in each quadrant. Color scale depicts gene expression in shWhite (log10 CPM; 2 replicate average). (C) Heatmaps of H2Av ChIP signal distribution at genes that are upregulated, downregulated or unchanged upon H2Av GLKD. Color scale depicts log2(IP/input). (D) Metaplot of the H2Av ChIP-seq data displayed in (C). (E) H2Av enrichment at the 1kb region flanking the TSS at all expressed genes, categorized based on differential expression upon H2Av GLKD.

## Supplemental Table Legends

**Table S1.** Primers

